# Genetic variation in the immunoglobulin heavy chain locus shapes the human antibody repertoire

**DOI:** 10.1101/2022.07.04.498729

**Authors:** Oscar L. Rodriguez, Yana Safonova, Catherine A. Silver, Kaitlyn Shields, William S. Gibson, Justin T. Kos, David Tieri, Hanzhong Ke, Katherine J. L. Jackson, Scott D. Boyd, Melissa L. Smith, Wayne A. Marasco, Corey T. Watson

## Abstract

Variation in the antibody response has been linked to differential outcomes in disease, and suboptimal vaccine and therapeutic responsiveness, the determinants of which have not been fully elucidated. Countering models that presume antibodies are generated largely by stochastic processes, we demonstrate that polymorphisms within the immunoglobulin heavy chain locus (IGH) significantly impact the naive and antigen-experienced antibody repertoire, indicating that genetics predisposes individuals to mount qualitatively and quantitatively different antibody responses. We pair recently developed long-read genomic sequencing methods with antibody repertoire profiling to comprehensively resolve IGH genetic variation, including novel structural variants, single nucleotide variants, and genes and alleles. We show that IGH germline variants determine the presence and frequency of antibody genes in the expressed repertoire, including those enriched in functional elements linked to V(D)J recombination, and overlapping disease-associated variants. These results illuminate the power of leveraging IGH genetics to better understand the regulation, function and dynamics of the antibody response in disease.

## Introduction

Antibodies (Abs) are critical to the function of the adaptive immune system, and have evolved to be one of the most diverse protein families in the human body, providing essential protection against foreign pathogens. The circulating Ab repertoire is composed of hundreds of millions of unique Abs^1, 2^, the composition of which varies considerably between individuals^1–3^, potentially explaining the varied Ab responses observed in a variety of disease contexts, including infection^4–8^, autoimmunity^9–12^, and cancer^13–15^. The initial formation of and diversity found within the Ab repertoire is mediated by complex molecular processes, and can be influenced by factors such as prior vaccination and infection, health status, sex, age, and genetics^16–21^. Delineating the mechanisms that drive variation in the functional Ab response is critical not only to understanding B cell-mediated immunity in disease, but also ultimately informing the design of improved vaccines and therapies^22, 23^. With respect to genetic factors, the impact of variants in the immunoglobulin heavy (IGH) and light chain loci on the antibody response has not been determined.

The human IGH locus is located immediately adjacent to the telomere of chromosome 14, and harbors 129 variable (V), 27 diversity (D) and 9 joining (J) genes that are utilized during V(D)J recombination to produce the heavy chain of an Ab^24^. The IGH locus is now understood to be among the most polymorphic and complex regions of the human genome^3, 25–29^. Akin to the extensive genetic diversity observed in the human leukocyte antigen (HLA) locus (>2,000 alleles), >680 IGH alleles have been cataloged solely from limited surveys^30^. In addition, IGH is highly enriched for large structural variants (SVs), including insertions, deletions, and duplications of functional genes, many of which show considerable variability between human populations^25, 29^. This extensive haplotype diversity and locus structural complexity has made IGH haplotype characterization challenging using standard high-throughput approaches, and as a result it has been largely ignored by genome-wide studies^25, 28, 31^. This has hindered our ability to assess the contribution of IGH polymorphism in disease phenotypes, and more fundamentally, our ability to conduct functional/molecular studies. We currently understand little about the genetic factors, and thus the associated molecular mechanisms, that dictate the regulation of the human Ab response. In fact, much of what we understand about the specific genomic factors involved in Ab repertoire development and variability comes from inbred animal models^32–35^, even though such questions would have greater relevance to health if addressed in outbred human populations^22^. These limitations continue to impede our understanding of the contribution of IGH polymorphism to disease risk, infection and response to vaccines and therapeutics^22, 31, 36, 37^.

Several lines of evidence now support the importance of IGH genetic variation in human B cell-mediated immune responses. Studies in monozygotic (MZ) twins have shown that many Ab repertoire features are correlated within twin pairs in both naïve and antigen-experienced B cell subsets, indicating strong heritable factors underlying repertoire variability^20, 21, 38^. Other studies have demonstrated that specific SVs and IG coding and regulatory element polymorphisms contribute to inter-individual variability in expressed human Ab repertoires^23, 39–42^. These observations, alongside biases in IG gene usage in various disease contexts, underscore potential connections between the germline and Ab function^39, 41, 43, 44^. Importantly, in many cases, key functional amino acids identified in disease-associated and antigen-specific Abs are encoded by polymorphic positions with variable allele frequencies among populations^23, 41^. These observations indicate that IGH variants could offer direct translational opportunities, with the ability to subset the population according to IG genotypes for more tailored healthcare decisions ^22^. However, investigations of the direct functional effects of human IGH germline variation conducted to date have been limited to only a small fraction of IGH variants known^39–42^.

In order to identify IGH polymorphisms that affect variation in the expressed Ab repertoire, we performed long-read sequencing to comprehensively genotype IGH, and combined these data with adaptive immune receptor repertoire sequencing (AIRR-seq) in 154 healthy adult individuals. We detected an extensive number of SNVs, indels and SVs across IGH, including novel IGH genes and alleles, and SVs collectively spanning >500 Kb. Using the AIRR-seq data to profile the IgM and IgG repertoire, we directly tested for effects of IGH variants on IGHV, IGHD and IGHJ gene usage frequencies. We show that for the majority of genes in the IgM and IgG repertoires, usage is associated with IGH germline polymorphism. Strikingly, for a subset of genes, IGH variants alone explain a large fraction of usage variation across individuals, and are strongly linked to IGH coding region changes. Finally, we found that IGH gene usage variants were enriched in regulatory elements involved in V(D)J recombination and overlapped SNVs previously associated to human phenotypes, offering insight into the underlying mechanisms linking germline variants to gene usage, and highlighting potential pathways from disease risk variant to phenotype. Our results clearly demonstrate that genetics plays a significant role in shaping an individual’s Ab repertoire, which will be critical to understand further in the context of human disease prevention and Ab-mediated immunity.

## Results

### Paired IGH targeted long-read and antibody repertoire sequencing

In this study, we compiled a dataset consisting of newly and previously generated germline IGH locus long-read sequencing data and AIRR-seq datasets^18^ in 154 healthy individuals (**Supplementary Table 1**). To our knowledge, this dataset represents the most comprehensive collection of matched full-locus IGH germline genotypes and expressed Ab repertoires. Samples in the cohort ranged in age from 17 to 78 years, and included individuals who self-reported as White (n=81), South Asian (n=20), Black or African American (n=19), Hispanic or Latino (n=19), East Asian (n=11), Native Hawaiian or Other Pacific Islander (n=1), American Indian or Alaska Native (n=1), or unknown (n=2).

Using our previously published method^28^, we performed probe-based targeted capture and long-read single molecule, real-time (SMRT) sequencing (Table 1; Supplementary Figure 1a,b) of the IGHV, D, and J gene regions (collectively referred to as IGH), spanning roughly ∼1.1 Mb from *IGHJ6* to the telomeric end of chromosome 14 (excluding the telomere). DNA used for each sample was isolated from either peripheral blood mononucleocytes (PBMCs) or polymorphonuclear leukocytes (PMNs). The mean coverage across IGH for all individuals ranged from 2X to 331X (mean=76X) with a mean read length ranging from 3.5 Kbp to 8.9 Kbp (mean=6.4 Kbp; Supplementary Figure 1c,d). Similar to our previously published work^28^, HiFi reads were aligned to a custom linear IGH reference inclusive of previously resolved insertions and used to generate local haplotype resolved assemblies. The mean assembly size across the total dataset was 2.3 Mb (range = .8 - 3.3 Mb), close to the expected diploid size of IGH (∼2.2 Mb), although the number and lengths of assembly contigs varied between platforms (Supplementary Figure 1e-g). These assemblies were then used to curate IGH gene/allele and variant genotype datasets (see below).

**Table 1.**
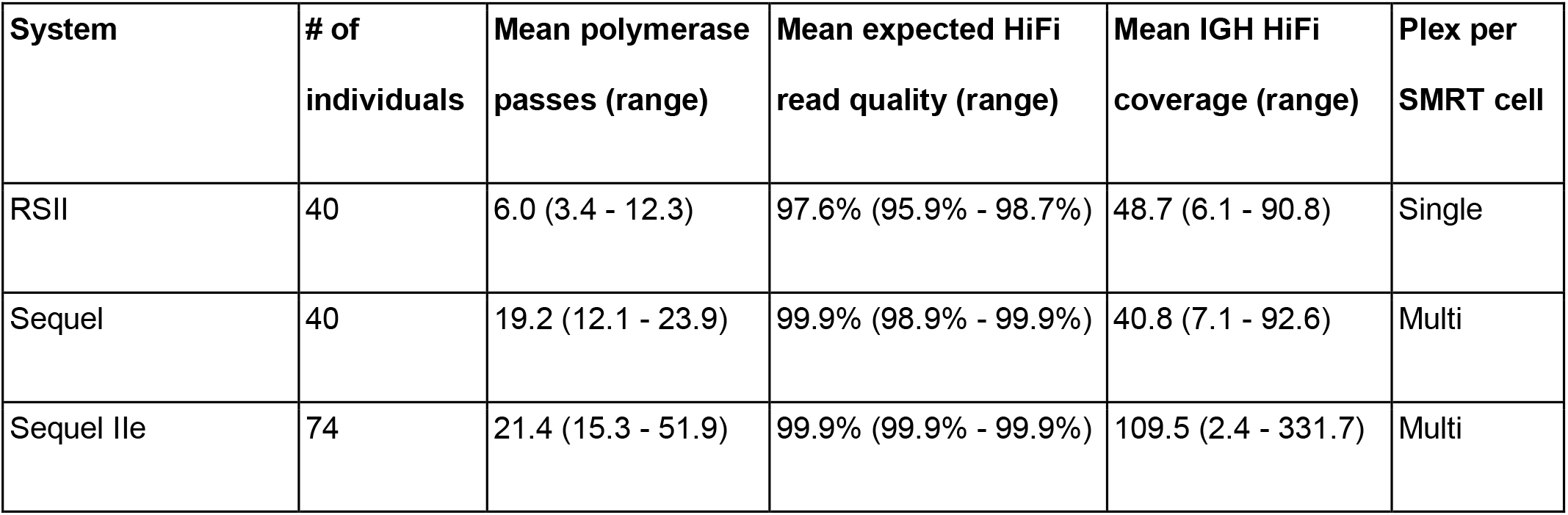
Sequencing statistics across SMRT sequencing systems.

AIRR-seq data was generated for 51 individuals using cDNA derived from total RNA isolated from PBMCs and sequenced using 5’ rapid amplification of complementary DNA ends (5’ RACE). For the remaining 103 individuals, previously generated PBMC derived AIRR-seq data for IgM and IgG was utilized. A standardized workflow was developed to process datasets generated using different protocols and sequencing methods^18^ (Methods). After processing, a mean of 9,038 B cell clones per repertoire was identified (Supplementary Figure 2a,b). The frequency of V, D and J genes within B cell clones was calculated (i.e., gene usage after collapsing sequences by clone) for each individual. Together these datasets allowed us to resolve large SVs and other genetic variants, and perform genetic association analysis with variation observed in the expressed Ab repertoire.

### Identification of large breakpoint resolved structural variants

A major goal of this study was to generate a high-confidence set of genetic variants and gene alleles in IGH in order to perform downstream genetic Ab repertoire association analysis (Fig. 1a). Previous reports have demonstrated that SVs are common in IGH, resulting in large insertions, deletions, duplications and complex events^25, 27–29, 45^. The presence of unresolved SVs can impact the accuracy of variant detection and genotyping. Thus, a key first step in the creation of genotype call sets was to breakpoint resolve and genotype SVs, which allowed us to account for SVs in determining homozygous, heterozygous, and hemizygous genotypes across all surveyed variants in the locus.

**Figure 1.**
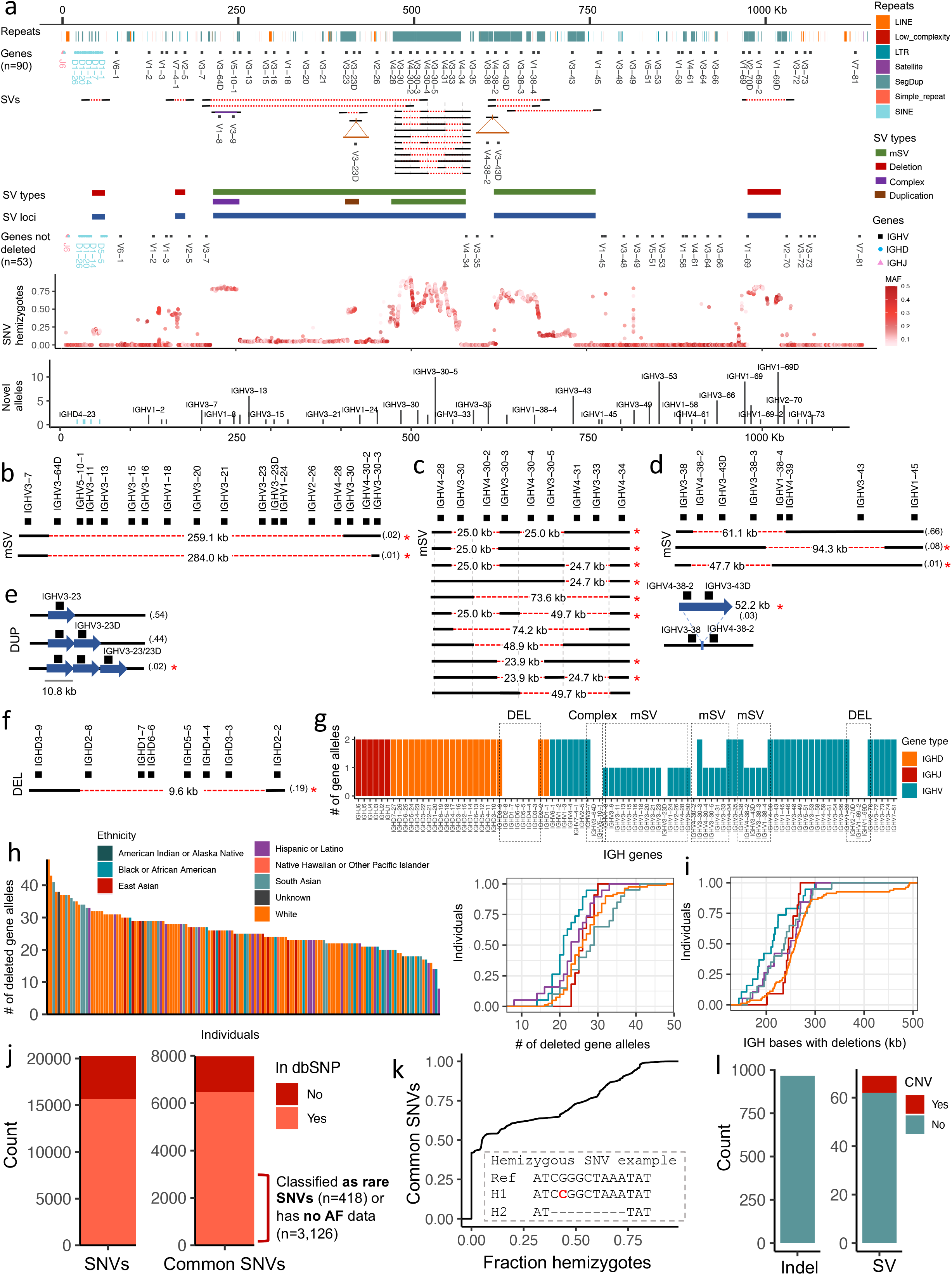
IGH genetic variation identified by long-read sequencing in a cohort of 154 individuals. **(a)** Map of the IGH locus with annotation tracks shown in the following order: repetitive sequences, joining (J), diversity (D) and variable (V) genes, structural variants (SV) resolved in this study, SV types, IGH loci with SVs, genes not deleted by SVs, fraction of hemizygotes across all common single nucleotide variants (SNVs), and number of novel alleles per gene. **(b)** A multi-allelic structural variant (mSV) with three alleles, including the reference assembly allele. Two of the SV alleles represent 259.1 and 284.9 Kbp deletions, deleting up to 16 genes. **(c)** mSV with 12 SV alleles. **(d)** mSV with 4 SV alleles: 3 deletions and 1 insertion representing a partial duplication relative to the reference. **(E)** Duplication SV including SV alleles harboring 1 to 3 copies of the *IGHV3-23* gene. Red asterisks (**b-e**) indicate SV alleles that were not previously resolved at the genomic level. **(f)** Deletion in the IGHD gene region that deletes 6 IGHD genes. **(g)** Count of alleles (n=36) deleted in an individual carrying multiple homozygous and hemizygous deletions. **(h)** Plots showing the number of gene alleles deleted for every individual in the cohort color coded by self-reported ethnicity and the CDF of deleted gene alleles per population. **(i)** Plot showing the CDF of the number of deleted bases in IGH associated with SVs, split by population. **(j)** Number of SNVs and common SNVs identified in the study cohort compared to the SNVs in dbSNP. A large portion (48%) of common SNVs identified here using long-read sequencing were missing, identified as rare, or had no allele frequency data in dbSNP. **(k)** The fraction of hemizygotes across all common SNVs. The embedded panel is an example of a hemizygous SNV. **(l)** The total count of indels and SVs identified.

Using a combination of haplotype-resolved assemblies and HiFi read coverage (Supplementary Figure 3a,b), we genotyped 8 large SV regions with SV alleles ranging in size from 9 Kbp to 284 Kbp (Fig. 1a-g; Supplementary Figure 4). These included deletions (n=3), a complex SV (n=1), a duplication (n=1) and multi-allelic SVs (mSV; n=3), two of which represented SV hotspots defined by > 2 SV alleles (Supplementary Table 2). Similar to other genetic variant types (e.g. SNVs) an SV allele is defined as an alternative sequence/haplotype relative to the reference. Of the 8 SV regions, the genomic positions for 3 overlapped. The three mSVs contained 3, 5 and 12 SV alleles and the duplication contained 3 SV alleles. In addition to the SV alleles described in Watson et al^25^,14 new SV alleles were breakpoint resolved, many of which are supported by previous AIRR-seq analysis ^26, 27, 46^. Detailed descriptions of these SVs are provided in the Supplementary Material.

The SV allele frequencies ranged from 0.01 to 0.73. On average across our cohort, relative to the reference assembly used in our analysis, we found that each individual carried 5.5 large SVs, resulting in the complete loss of 6.7 genes (range = 0 - 17), 26.11 gene alleles (range = 14 - 48), and deleted diploid bases summing to 257 Kbp of the locus (range = 49 - 493 Kbp; Fig. 1h,i). The observed number of genes and bases deleted within individuals varied by self-reported ethnicity (Fig. 1i). In total, 31 out of 54 IGHV and 6 out of 26 IGHD genes were removed by 1 or more of the SVs identified in at least one sample (Fig. 1a).

### Long-read sequencing identifies SNVs, indels and smaller SVs within IGH

SNVs and indels are difficult to characterize within segmental duplications and SVs. Here, we used haplotype-resolved assemblies to more accurately detect and genotype SNVs. In total we identified 20,510 SNVs in one or more individuals, of which 7,980 (39%) were common, defined by a minor allele frequency (MAF) => 0.05 (Fig. 1j). While the majority (97%) of all non-redundant SNVs were in non-coding regions, 472, 103 and 40 SNVs were within exons, introns and recombination signal sequences (RSS), respectively. Interestingly, SNVs within these genomic features were non-uniformly distributed across IGHV genes (Supplementary Figure 5). For example, while the mean number of SNVs in IGHV gene RSS was 0.68, several genes, including *IGHV3-21* and *IGHV3-66* had 7 and 5 SNVs in their RSS, respectively. Similarly, the mean number of SNVs across IGHV introns was 1.7, but *IGHV3-23*, *IGHV4-39* and *IGHV7-81* had 9, 8 and 8 intronic SNVs, respectively.

Based on earlier reports of elevated numbers of SNVs in the IGH locus^25^, we hypothesized that many of the SNVs identified in this cohort would be novel. Indeed, a total of 4,625 (23%) SNVs had not been previously identified cataloged in dbSNP (release 153), including 1,513 (19%) common SNVs (Fig. 1j). Of the total SNVs not in dbSNP, 2,393 (59%) were within SVs. Even though a large portion of common SNVs were in dbSNP, we found that 3,126 (48%) of the common SNVs had no allele frequency data and 418 (6%) were labeled as rare variants (Fig. 1j). Thus in total, 63% (5,057) of common SNVs identified in our cohort were either missing from dbSNP or are lacking accurate genotype information.

The incomplete and inaccurate genotype frequency information available in dbSNP for IGH is likely in part caused by the prevalence of large SVs in the region, which have hindered the analysis of standard high-throughput genotyping approaches. This is supported directly in our data, as 3,406 (43%) of the common SNVs we identified reside within SVs. Here, since SNVs were detected by aligning both haplotype assemblies to the reference, SNVs overlapping heterozygous deletions were simultaneously detected and genotyped as hemizygous. Hemizygous SNVs are often genotyped as homozygous when using short-read and/or microarray data and are excluded from studies due to a departure from Mendelian inheritance and Hardy-Weinberg equilibrium^47^. We observed that the frequency of hemizygous individuals was greater at 2,136 (27%) common SNVs than individuals with both chromosomes present (Fig. 1a,k). Critically, analysis of SNVs within the complex SVs we identified was possible due to long-read assemblies, highlighting the utility of long-read data in IGH beyond assembly and SV detection.

In addition to SNVs and large SVs, we identified indels (2-49 bp) and small non-coding SVs (50 bp - 9 Kbp) using haplotype-resolved assemblies and validated these using mapped HiFi reads (Fig. 1l). In total, 966 indels and 71 small SVs were detected, including expansions and contractions of tandem repeats, mobile element insertions and complex events. We additionally observed highly polymorphic indels and SVs (Supplementary Figure 6). For example, a tandem repeat with a motif length of 86 bp 5 Kbp upstream of *IGHV3-20* contained 7 tandem repeat alleles ranging in motif copies from 3 to 9 (Supplementary Figure 6a). Another example includes a complex SV between *IGHV1-2* and *IGHV1-3* with three SV alleles containing multiple copies of a tandem repeat with low sequence matches between motif copies (Supplementary Figure 6b). An alignment between the 3 SV alleles contains multiple mismatches including base differences, insertions and deletions.

### Identification of novel IGH gene alleles using long-read sequencing

Analysis of AIRR-seq data critically relies on the assignment of AIRR-seq reads to specific IGHV, D, and J gene alleles using existing germline databases. Accurate assignments of reads to gene alleles is used for analyzing a variety of Ab repertoire features including gene usage and somatic hypermutation. In order to obtain a more complete allele database, we used haplotype-resolved assemblies to annotate additional undocumented novel alleles, defined as alleles absent from the ImMunoGeneTics Information System (IMGT; imgt.org) germline database. In total, we identified 125 IGHV and 5 IGHD high-confidence putative novel alleles, conservatively defined as alleles with exact matches to 10 or more HiFi reads, or identified in two or more individuals (Supplementary Table 3). Of these 125 IGHV alleles, 72 (58%) were found in at least 2 individuals; 23 (18%) and 9 (7%) were found in at least 5 and 10 individuals, respectively (Fig. 1a); the remaining 53 alleles were found in only one sample, but were supported by >=10 HiFi reads. Of the 5 novel IGHD alleles, 4 were found in at least 2 individuals and 3 were found in 14 or more individuals. In total, the discovery of 125 and 5 novel IGHV and IGHD alleles represents a 37% and 11% increase in the number of IMGT-documented IGHV and IGHD F/ORF alleles, respectively.

### Gene usage in the expressed antibody repertoire is strongly associated with common IGH variants

Across the genome, genetic variation has consistently been associated with molecular phenotypes such as gene expression and splicing^48^. Performing such analysis on repetitive and SV dense loci such as IGH has been limited by the use of short-read or microarray derived variants. Here, in order to determine if the long-read sequencing derived genetic variants described above impact the expressed Ab repertoire, we used a quantitative trait locus (QTL) framework (see Materials and Methods) to test if gene usage in the naive (IgM) and antigen-experienced (IgG) repertoire was associated with variant genotypes. The clonal gene usage in 50, 25 and 6 IGHV, IGHD and IGHJ genes, respectively, was tested against all common genetic variants (7,042 SNVs, 223 indels, 32 SVs) including SV alleles at 6 of the 8 large (> 9 Kbp) SV regions. In the IgM repertoire, after stringent multiple-testing correction (Bonferroni), 4,025 variants (3,967 SNVs, 50 indels and 8 SVs) were associated with usage of 37 (74%), 20 (80%) and 2 (33%) IGHV, IGHD and IGHJ genes, respectively (Fig. 2a). Similar results were observed in the IgG repertoire: 3,675 variants (3,628 SNVs, 36 indels and 11 SVs) were associated with gene usage variation in 33, 14, and 3 IGHV, IGHD and IGHJ genes, respectively (Supplementary Figure 7). Of those genes, all but 3 and 2 IGHV and IGHJ genes, respectively, overlapped those observed in the IgM repertoire (Supplementary Figure 8), and were associated with 3,320 genetic variants in both repertoires, providing evidence that genetic effects impacting the naive repertoire extend to the antigen-experienced repertoire. The relationship between IgM and IgG gene usage is further demonstrated by the significant (*P value* < 0.05) gene usage correlation between both repertoires (Supplementary Figure 8c).

**Figure 2.**
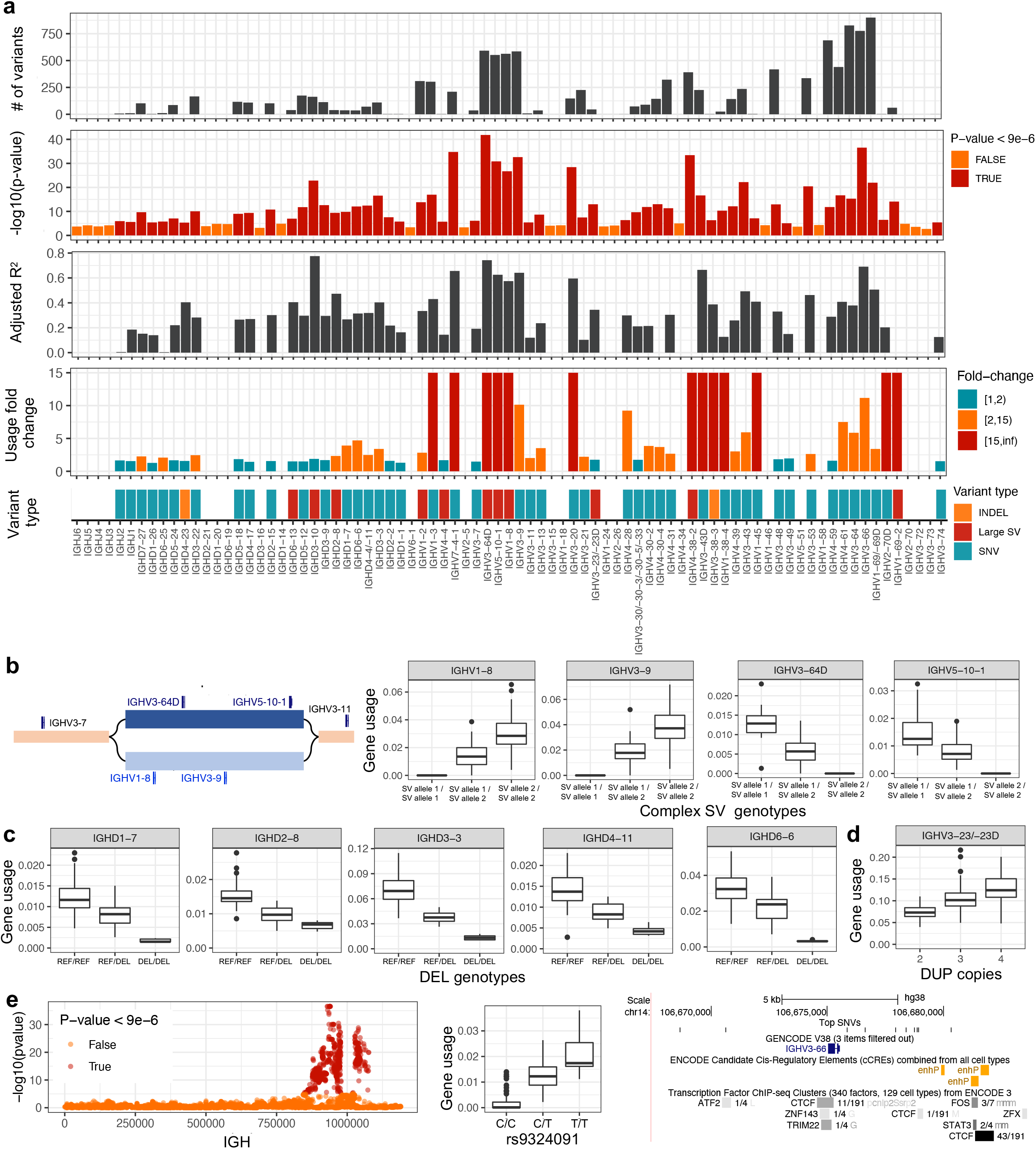
IGH variants have significant impact on gene usage in the IgM repertoire. **(a)** Per gene statistics from guQTL analysis in the IgM repertoire, including: (i) the number of associated variants (*P value* < 9e-6); the (ii) *P value*, (iii) adjusted R^2^ for variance in gene usage explained, (iv) the fold change between genotypes, and (v) the variant type for the lead guQTL variant. **(b)** Gene usage guQTL results for the genes *IGHV1-8*, *IGHV3-9*, *IGHV3-64D* and *IGHV5-10-1*. The genomic copy number and usage of these genes is associated with a complex SV, shown as a genome graph. The SV alleles (light and dark blue bars) contain unique sequences and are mutually exclusive. Individuals homozygous for the SV allele with *IGHV3-64D* and *IGHV5-10-1* (genotype group “SV allele 1/SV allele 1”) have higher usage frequency of those genes than individuals heterozygous or homozygous for the alternate SV allele. **(c)** Gene usage for genes within the IGHD gene region deletion. Individuals homozygous for the deletion (genotype group “DEL/DEL”) use those genes at lower frequency than the rest of the cohort. **(d)** Gene usage for *IGH3-23/-23D* between individuals with varying gene copy numbers. As expected, individuals carrying more gene copies use these genes at higher frequencies. **(e,f)** SNVs associated with the usage of genes **(e)** *IGHV3-66* and **(f)** *IGHV1-2.* The Manhattan plot shows the - log10(p-value) for all SNVs in the IGH locus tested for each gene. Dark red SNVs are those that passed Bonferroni correction (p-val < 9e-6). Usage between the SNV genotypes for the most significant SNV is shown along with the genomic localization of the lead SNVs. For *IGHV3-66,* there are 10 lead SNVs with the same p-value.

Thus, collectively, across the two repertoires, 4,380 unique variants (4,310 SNVs, 58 indels and 12 SVs) were associated with gene usage changes in 40 (80%), 20 (80%), and 4 (66%) unique IGHV, IGHD and IGHJ genes. Summary data for each gene analyzed in our dataset is provided in Supplementary Table 4 for IgM and IgG. This includes: (1) the number of significant gene usage QTL (guQTL) variants identified; (2) the significance level of the lead guQTL (-log10 *P value*), defined as the variant with the lowest *P value*; (3) lead guQTL variant type (SNV, indel, SV); (4) the variance explained by the lead guQTL; and (5) the mean fold change in usage between the reference and alternate genotypes. Given the gene usage correlation and high guQTL overlap between IgM and IgG, and the fact that gene usage is a product of V(D)J recombination, we focus on the IgM repertoire in the following results sections.

Given the extent of SVs that alter gene copy number within IGH, we expected to observe significant effects of large SVs on gene usage. Indeed, within the IgM repertoire, there were 5 IGHD genes and 6 IGHV genes that resided within SV regions, and for which the lead guQTL variant was the SV itself or a variant in high LD with the SV (r > .9; Fig. 2a). These SV associations were among the most statistically significant in this dataset, and explained between ∼20% and >77% of the variation in IgM usage observed for associated genes (Fig. 2a). The most significant association identified was for *IGHV3-64D* (*P value* =1.46E-42; Fig. 2a), involving a complex SV, which alters the genomic copy number of 4 functional IGHV genes (*IGHV3-64D*, *IGHV5-10-1*, *IGHV1-8*, and *IGHV3-9*) from 0-2 diploid copies (Fig. 1a). The impact on gene usage of this SV was as expected, following an additive model in which individuals with zero copies of a given gene had the lowest mean usage (in this case 0%), whereas individuals with 2 diploid copies of a given gene had the highest mean usage, and heterozygotes showed intermediate usage (Fig. 2b). Other large deletions followed a similar pattern. For example, the deletion spanning the genes *IGHD2-8* to *IGHD3-3* was significantly associated with the usage of six IGHD genes (Fig. 1f), five of which reside within the deletion *(IGHD2-8*, *IGHD1-7*, *IGHD6-6*, *IGHD4-11/4-4* and *IGHD3-3*; Fig. 2c); these results were consistent with those noted previously^42^. Due to sample size (n=7), the largest mSV deleting 16 IGHV genes was not tested; however, we observed empirically that individuals carrying either one of these large deletions had decreased usage across 15 out of the 16 genes (Supplementary Figure 9). In addition to SVs that resulted in gene deletions, we also noted a significant association with the duplication characterized for the *IGHV3-23/D* genes, at which we tested for effects of copy number genotypes between 2 to 4 diploid copies (Fig. 2d). Again, this effect was consistent with an additive contribution of gene copy number, with mean usage increasing incrementally from 7.4% in individuals with 2 copies, to ∼13% in individuals with 4 copies; individuals carrying the rare 3-copy haplotype (Fig. 1e) were excluded from this analysis (Supplementary Figure 10).

We additionally identified 3 IGHD genes (*IGHD6-13*, *IGHD3-9* and *IGHD3-10*) and 2 IGHV genes (*IGHV1-2 and IGHV4-4*) that were most significantly associated with SVs or a variant in high linkage disequilibrium (LD, r^2^ > .9) with a SV, although the copy number of these genes was not directly altered. The deletion spanning the IGHD genes mentioned above was the most significant variant associated with *IGHD3-10* usage, even though the gene is ∼3 Kbp away from the deletion. Contrary to genes residing within the deletion, the mean usage of *IGHD3-10* increased from 10% to 19% in individuals with the deletion on both haplotypes (Supplementary Figure 11), suggesting that the deletion modulated the usage of these genes through cis-regulatory mechanisms^49, 50^. Interestingly, usage of the gene, *IGHV1-69-2*, which resides within a deletion SV, was most significantly associated with a secondary SV, located ∼322 Kb away. However, given the low usage of *IGHV1-69-2,* deeper repertoire sequencing will likely be needed to tease out the effect of both SVs.

We next focused on the 42 genes (IGHJ, n=2; IGHD, n=12; IGHV, n=28) that were not significantly associated with large SVs. The lead guQTLs associated with 40 of these genes were SNVs, and the remaining 2 were indels; although we identified the presence of smaller SVs and tandem repeats in our dataset, none of these were found to be lead variants in our analysis. For 38 of the genes, we identified from 2 to 900 guQTLs, reflecting local haplotype structure (Fig. 2a). In some cases, a SNV or indel was the lead guQTL for genes residing within SVs indicating that multiple variant types need to be taken into account to fully model the genetic effects on usage (see below). Similar to SVs, the lead guQTL SNVs/indels explained a significant fraction of usage variation, in some cases up to 68% (range, R^2^ = 0.003 - 0.69; mean = 0.29), exhibiting large usage differences between genotype groups (Fig. 2a). The lead guQTLs for all 42 genes resided within non-coding regions. The median genomic distance between intergenic guQTLs and genes was 5.1 Kbp (min = 13 bp, max = 1.1 Mbp).

The most significant SNV-driven guQTL in this dataset was for *IGHV3-66* (*P value* = 2.86e-37; Fig. 2a). In total, there were 776 SNVs associated with the usage of *IGHV3-66* (Fig. 2a,e). These included 10 lead SNVs (*r*^2^ =1), spanning a region of 11.6 Kbp surrounding the gene, which explained ∼69% of variation in usage, representing a mean fold-change in usage of 11.2-fold between the two homozygous genotypes (Fig. 2a,e).

### Conditional analysis identifies multiple variants associated with the usage of single genes

Previous eQTL studies have demonstrated that multiple independent variants can influence gene expression^48^. Here, we hypothesized that the usage of individual genes could be affected by multiple variants, such as multiple SNVs, or a combination of variant types. To test this, we performed a conditional analysis by running an additional QTL analysis in individuals homozygous for either the reference or alternate allele for the lead guQTL variant of all significantly associated genes. Out of the 59 genes associated with gene usage in the IgM repertoire, 55 genes were tested for additional associations. The 4 genes not tested had fewer than 50 individuals with homozygous reference or alternate allele genotypes. From this analysis, we identified an additional variant associated with the usage of 14 genes (Supplementary Table 5). In combination with the initial guQTL defined above, for 12 of these 14 genes, we observed effects of 2 SNVs, and in the remaining, we observed combined effects of an SV and SNV. The mean genomic distance between the lead and secondary guQTL variants was 36.2 Kbp (range = 1.7 - 161.4 Kbp). Here, we present *IGHV1-2* and *IGHV3-66* as examples of genes associated with 2 independent variants. Data for all genes is provided in Supplementary Table 5.

For *IGHV1-2*, the lead guQTL was a SV ∼31 Kb away from *IGHV1-2* (Fig. 3a), which involved the deletion of *IGHV7-4-1.* Individuals homozygous for the deletion used *IGHV1-2* at a 2.8-fold higher rate than individuals homozygous for the SV insertion allele (Fig. 3a). Conditioning on individuals without the deletion identified 35 SNVs additionally associated with the usage of *IGHV1-2* (Fig. 3b). Of these individuals, heterozygotes for the lead conditional guQTL used *IGHV1-2* at a level (mean usage = 3.8%) similar to those with a deletion in both haplotypes (mean usage = 4.2%). Sequencing data from heterozygotes at the lead conditional guQTL were inspected manually to confirm that *IGHV7-4-1* deletions were not present in these individuals.

**Figure 3.**
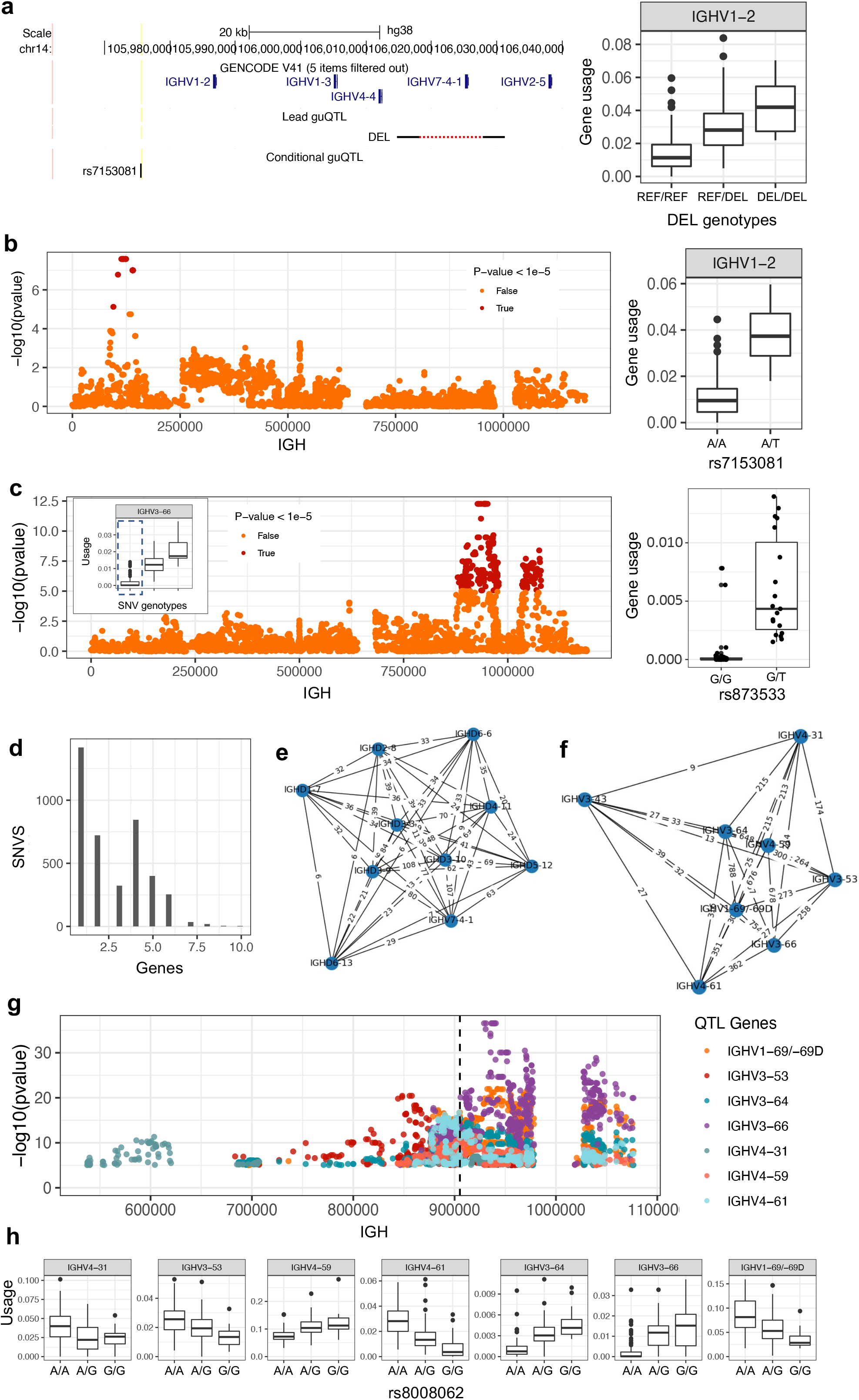
Conditional analysis and construction of an IGH guQTL network reveals coordinated genetic effects on gene usage patterns. **(a)** UCSC Genome Browser showing the lead and conditional guQTL for *IGHV1-2*. Individuals with more copies of the large deletion SV have greater *IGHV1-2* usage (right). **(b,c)** Conditional analysis for *IGHV1-2* and *IGHV3-66* finds additional variants associated with gene usage. Manhattan plots (left) show the statistical significance of all SNVs tested for secondary effects on gene usage (red indicates Bonferroni corrected significant SNVs), after selecting individuals from a single genotype group from the original lead guQTL. Inset boxplots show gene usage variation for each gene, partitioned by genotypes at the lead eQTL, and the Individuals selected for conditional analysis are indicated by the dashed box. Box plots (right) show gene usage variation partitioned by genotypes at the secondary guQTL. **(d)** Bar plot showing the number of genes associated with each SNV in the primary guQTL analysis (Fig. 2A). **(e,f)** Graphs of example cliques identified from a comprehensive network of genes and guQTL variants (Supplementary Fig. 12,13), demarcating groups of genes associated with overlapping sets of guQTLs. For each clique (**e, f**), genes are shown as nodes, connected by edges displaying the number of shared guQTLs. **(g,h)** Example of a single SNV (dotted line) associated with 7 IGHV genes with coordinated usage patterns. Manhattan plot **(g)** showing statistically significant SNVs (points) associated with the usage of 7 genes; each point is colored by the gene it is associated with. The position of an example SNV associated with all seven genes is indicated by the dashed line. Boxplots (**h**) show usage variation for each gene partitioned by genotypes at this SNV.

For *IGHV3-66*, the lead guQTL was a SNV. Individuals homozygous for the reference and alternate allele had a mean usage of 0.19% and 2.14%, respectively. By conditioning on this variant, considering only individuals homozygous for the reference allele, a total of 438 additional SNVs were significantly associated with *IGHV3-66* usage (Fig. 3c). At the most significant SNV from this analysis, only reference allele homozygotes and heterozygotes were observed. In heterozygotes, the mean usage was 0.006% compared to 0.0003% in homozygotes, with many individuals in the homozygote group exhibiting 0% usage (Fig. 3c). Thus, based on this conditional guQTL analysis, variation in *IGHV3-66* usage can be further explained even in individuals with relatively low usage.

### Gene by guQTL network analysis reveals that the usage of multiple genes is associated with overlapping sets of variants

In addition to discovering multiple variants associated with the usage of a single gene, our guQTL association analyses also identified single variants associated with the usage of multiple genes. This was intriguing as V(D)J recombination studies in animal models have demonstrated the coordinated selection of genes through the same regulatory elements^32, 51^. In mice, IG V genes reside in topologically associating domains (TADs) and disruption of regulatory elements within the IG loci has been shown to cause altered gene usage within these domains^52–54^. Given this, we further assessed coordinated genetic signals involving sets of multiple variants and genes. We found that 2,607 (66%) guQTL variants were associated with > 1 gene (Fig. 3d). We reasoned that this could have multiple underlying causes: (1) the SNV is tagging a SV overlapping multiple genes; (2) the SNV is tagging multiple causative regulatory SNVs; (3) the SNV is overlapping a regulatory element controlling multiple genes; or (4) a combination of any of the prior explanations.

To determine the set of guQTL genes with the same set of guQTL variants, we created a network with genes as nodes and edges connecting genes associated with the same guQTL SNVs (Fig. 3e; Supplementary Figure 12). The weight of the edges corresponded to the number of guQTL SNVs connecting two genes. A total of 23 cliques (subgraphs in which all genes are connected) were identified with edge weights greater > 2 (i.e., more than 2 SNVs connecting 2 genes). These 23 cliques included a total of 16 IGHD and 29 IGHV genes, with the number of genes per clique ranging from 2 to 9. Out of the 23 cliques, 10 were primarily composed of genes within SVs (Fig. 3e; Supplementary Figure 13).

We also identified cliques made up primarily of genes outside of SVs (Fig. 3f). For example, the SNV shown in Figs. 3g was associated with the usage of 7 genes, *IGHV4-31*, *IGHV3-53*, *IGHV4-59*, *IGHV4-61*, *IGHV3-64*, *IGHV3-66* and *IGHV1-69/-69D*; this variant was located ∼120 Kbp away from the nearest SV, and exhibited low LD with the SV (*r*^2^ = 0.09). Interestingly, gene usage patterns associated with this SNV were either negatively or positively correlated depending on the gene. Individuals homozygous for the reference allele had higher usage of *IGHV4-31, IGHV3-53*, *IGHV4-61* and *IGHV1-69/-69D* and lower usage for the remaining genes. In summary, we show that the usage of specific sets of genes in the repertoire are associated with the same sets of variants, indicating the potential for complex and coordinated regulatory mechanisms.

### Variants associated with gene usage variation are enriched in regulatory regions involved in V(D)J recombination

Large scale studies using expression, epigenomic and disease or trait-associated variant datasets have identified non-coding variants in regulatory elements linked to their phenotypes of interest ^48, 55–57^. Specific to V(D)J recombination, recombination signal sequences (RSS) are sequence motifs in IG and T cell receptor non-coding regions used by RAG1/RAG2 proteins to direct double-strand DNA breaks and initiate somatic recombination^58^. Additionally, CTCF and cohesin binding has been shown to regulate contraction and recombination of IGH^59–61^. We therefore hypothesized that variants might modulate gene usage through regulatory elements such as CTCF-binding sites. To test this, we tested for the enrichment of guQTL SNVs within ENCODE Registry candidate cis-Regulatory Elements (cCREs) (Fig. 4a). The cCRES were split into 9 classifications: (1) CTCF-only and CTCF-bound, (2) proximal enhancer-like and CTCF-bound, (3) proximal enhancer-like, (4) DNase and H3K4me3, (5) promoter-like, (6) distal enhancer-like, (7) distal enhancer-like and CTCF-bound, (8) DNase, H3K4me3, and CTCF-bound, and (9) promoter-like and CTCF-bound. Using a one-sided Fisher exact test, we determined that SNVs were significantly enriched within CTCF-only and CTCF-bound (Fishers exact, *P value* = 3.8e-04) and distal enhancer-like and CTCF-bound (*P value* = 0.014). DNAse and H3K4me3 cCRE was nominally significant (Fishers exact, *P value* = .08). A total of 23 out of 3,573 guQTL SNVs tested were within CTCF-only and CTF-bound cCRE compared to 2 out of 2,419 common non-guQTL SNVs. These 23 SNVs were significantly associated with 3 IGHD genes and 19 IGHV genes and resided within 12 distinct cCREs. Interestingly, 4 SNVs within a CTCF-only and CTCF-bound cCRE (ENCODEAccession: EH38E1747546; chr14:106695880-106696139 (hg38)) were found between *IGHV3-66* and *IGHV1-69* and associated with usage of *IGHV3-53, IGHV4-59*, *IGHV3-66*, *IGHV3-64* and *IGHV1-69/-69D*. These genes are also included in a clique associated with the same set of guQTL SNVs (Fig. 3e-g). Within the DNAse and H3K4me3 cCREs, there were 10 SNVs associated with gene usage for 8 and 2 IGHD and IGHV genes, respectively. H3K4me3 is critical for V(D)J recombination via interaction with RAG2; disruption of the binding between RAG2 and H3K4me3 has been shown *in vivo* to reduce V(D)J recombination^62^.

**Figure 4.**
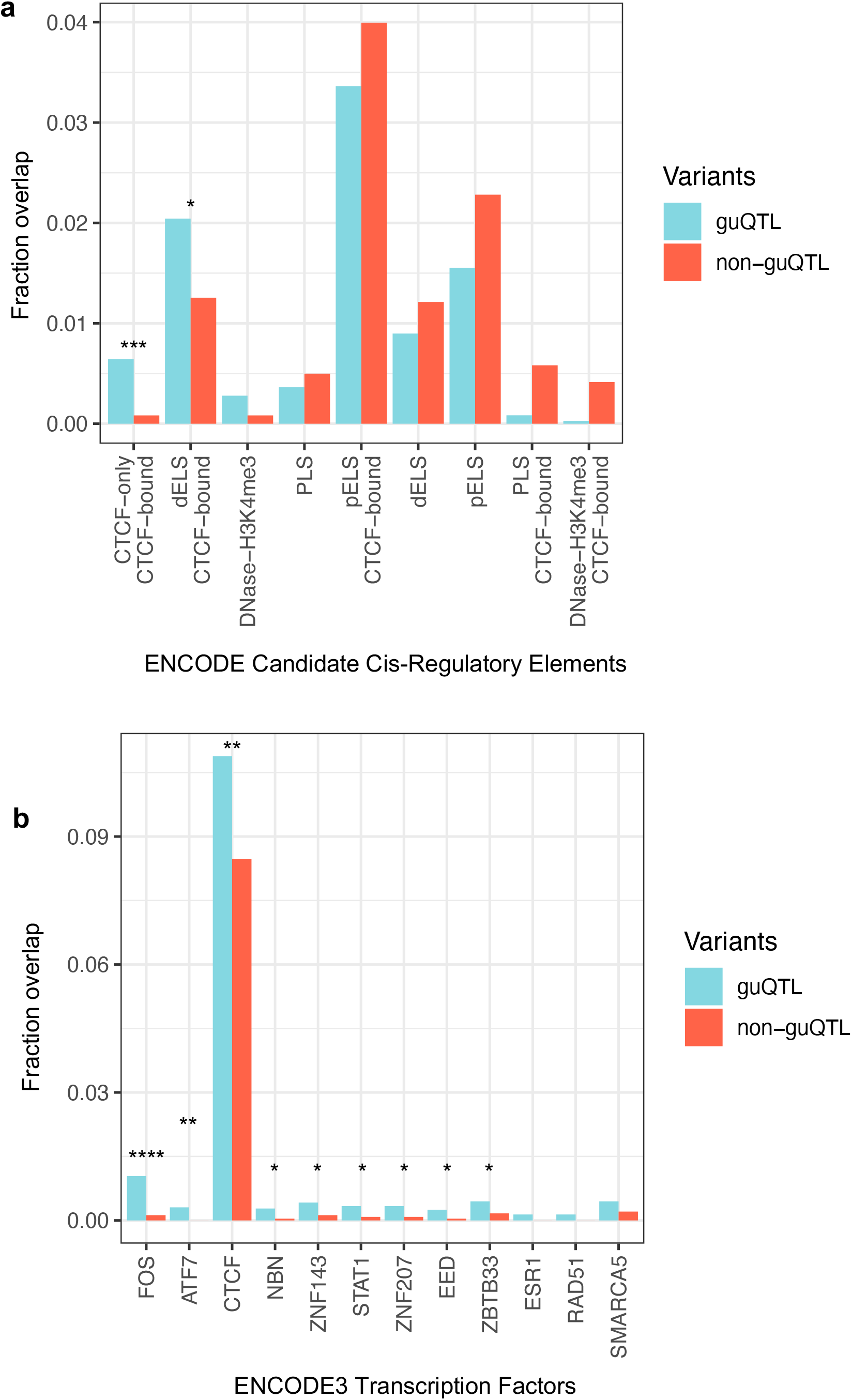
Enrichment of guQTL variants in regulatory elements and transcription factor binding sites involved in V(D)J recombination. **(a, b)** Bar plots showing the fraction of guQTL SNVs (“QTL”) that overlapped (**a**) ENCODE candidate cis-regulatory elements, and (**b**) ENCODE3 TFBS, compared to the overlap observed for the complete set of variants used in the guQTL analysis (“All”). Regulatory elements and TFBS for which statistically significant enrichments were observed are indicated by asterisks: Fisher’s Exact Test; * *P value* < 0.05; ** *P value* < 0.005; \*\*\**P value* < 0.0005; \*\*\*\**P value* < 0.00005;

We additionally compared the enrichment of guQTLs in specific transcription factor binding sites (TFBS) using the ENCODE3 Transcription Factor ChIP-seq binding site dataset (Fig. 4b). A total of 365 TFBS with high normalized ChIP-seq signals were tested. Again, an enrichment of guQTLs in the CTCF binding sites was observed (Fishers exact, *P value* = 0.004). Significant enrichments were observed for eight additional TFBSs (P *value* < 0.05) including EED.. Interestingly, the disruption of *Eed* in mice has been shown to affect IGHV gene usage^53^. The fact that SNVs are enriched in sites associated with V(D)J recombination rather than transcription (e.g. promoters and enhancers) is enticing, and provides strong support that the guQTLs identified here may impact gene usage via effects on V(D)J recombination.

### Genotypes within IGH coding regions and guQTLs are strongly associated

IGH germline coding variants can directly alter Ab function by modifying antigen binding^23, 63, 64^, and previous studies have demonstrated that specific coding alleles are utilized at different frequencies within the repertoire^23, 41^. To assess this more comprehensively in our dataset, we tested for associations between IGH gene alleles and all lead guQTLs (Supplementary Table 4). We found that allele frequency distributions at 21 IGHV genes were significantly different based on lead guQTL genotype (Fisher exact test *P value* < 0.05; Fig. 5a). The top three genes with the largest allele differences between guQTL variant genotype groups were *IGHV3-64* (*P value* = 6.9e-57; Fig. 5b), *IGHV3-53* (*P value* =4.4e-54; Fig. 5c), and *IGHV3-66* (*P value* = 5.0E-49; Fig. 5d). In the case of *IGHV3-66*, out of the 62 individuals who were homozygous for the reference allele at the lead *IGHV3-66* guQTL, 35 (52%) and 15 (23%) were homozygous and heterozygous, respectively, for the *IGHV3-66*03* allele. In contrast, *IGHV3-66*03* was not observed in any of the individuals homozygous for the alternate allele at this guQTL, which were all homozygous for *IGHV3-66*01*. These results show a direct genetic link between gene usage and coding variation, indicating that both should be considered in future studies investigating germline effects on Ab function.

**Figure 5.**
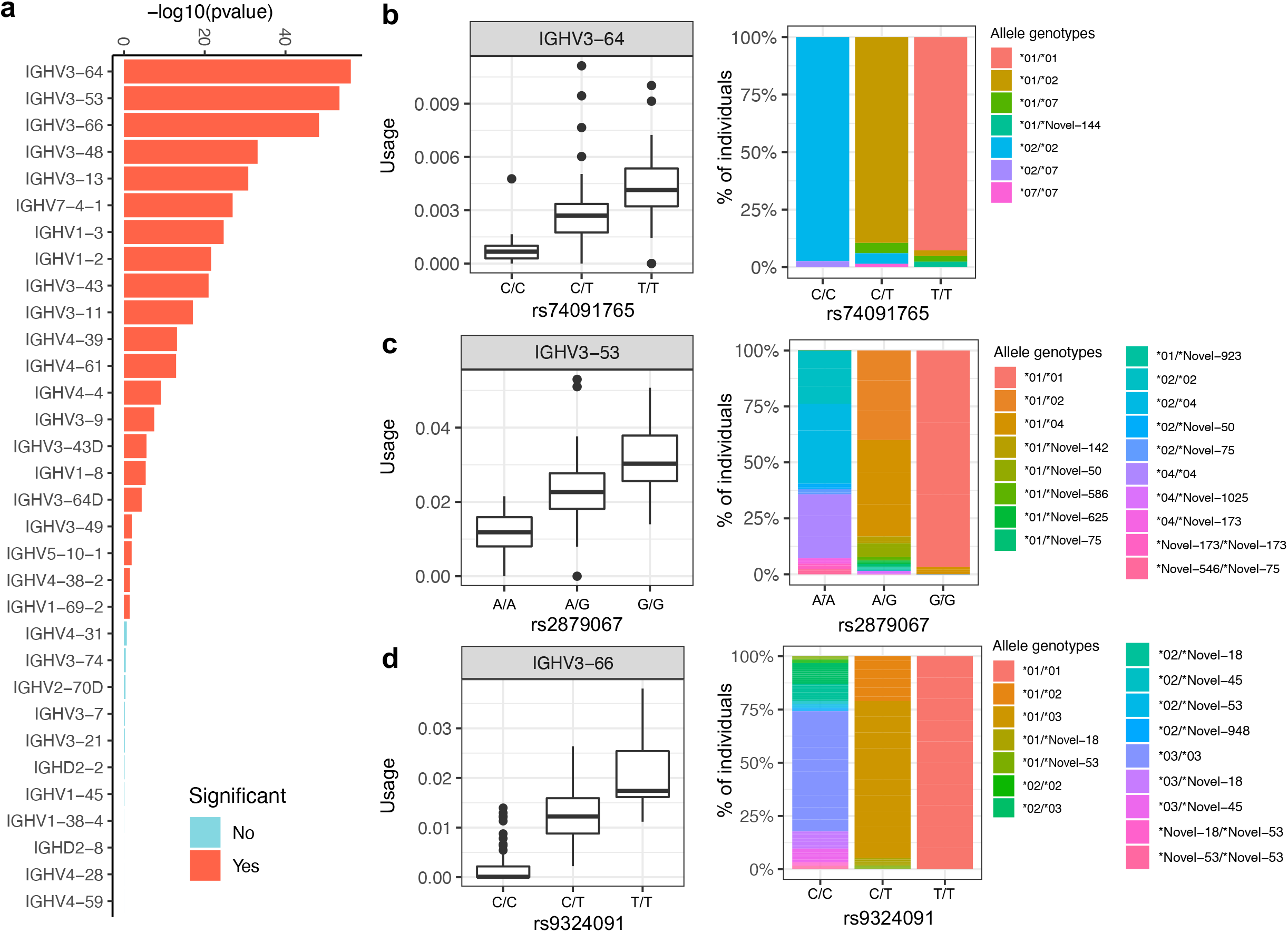
Association between IGHV coding region polymorphism and guQTL genotypes. **(a)** For each IGHV gene, differences in the distribution of coding region allele-level genotypes among individuals partitioned by genotype at the lead guQTL for that gene was assessed (Fisher’s exact test). Bar plot showing -log10(p-value) for each gene from this analysis; bars are colored based on statistical significance (*P value* < 0.05), red indicating genes for which coding allele genotype distributions were skewed based on guQTL genotype. **(b-d)** For the three most significant genes from this analysis (**a**), IgM gene usage (box plots) at the lead guQTL for each gene, and the distributions (stacked bar plots) of the respective coding allele genotypes, partitioned by guQTL genotype are provided. variant genotype group and the gene alleles genotypes in each guQTL variant genotype group is shown for *IGHV3-64*, *IGHV3-53* and *IGHV3-66*.

### GWAS disease risk and trait variants overlap guQTLs

Biased gene usage has consistently been observed in autoimmune and infectious diseases^37, 65^. We have argued that one possible explanation for these biases is that they are mediated through genetic variants that influence Ab antigen specificity and/or gene usage^22^. Integrating genome-wide association (GWAS) and eQTL datasets has been an effective method for assessing the potential function of risk variants to better understand links between genetic variation and disease pathology^48, 66, 67^. Here, we assessed whether IgM and IgG guQTL SNVs were also identified by GWASs (Fig. 6a). In total, across IGH (chr14:105,860,000-107,043,718, GRCh38) there were 41 SNVs associated with 17 traits/diseases reported in the NHGRI GWAS catalog (*P value* > 4-e6). In total, 22 SNVs from 10 independent GWAS performed on 8 diseases/traits overlapped guQTL SNVs. These included SNVs associated with rheumatic heart disease (RHD) and Kawasaki disease (KD). In both diseases, SNVs were significantly associated with the usage of genes previously implicated by GWAS (*IGHV4-61* for RHD and *IGHV3-66* for KD)^63, 68^. In the case of RHD, the risk variant identified in IGH is the strongest genetic association identified to date^63^; this association implicated *IGHV4-61*02* in increased risk. Interestingly, only individuals with the GWAS-guQTL SNV reference allele carried *IGHV4-61*02*, and these individuals had significantly lower *IGHV4-61* usage in IgM and IgG. In both RHD and KD, the usage of additional genes were also associated with the same guQTL SNV. For KD, the SNVs detected in the GWAS were also associated with *IGHV1-69/-69D*, *IGHV3-64* and *IGHV4-61* usage (Fig. 6b). Similar to using expression data to prioritize genes affected by SNVs identified from GWAS, here we show that guQTL-GWAS SNVs are associated with the usage of multiple genes in the Ab repertoire. Additional diseases/traits associated with SNVs identified by both GWAS and our guQTL analysis included the proportion of morphologically activated microglia in the midfrontal cortex, and estradiol levels, which were associated with the usage of *IGHV1-69/-69D* and *IGHV2-70D*, and *IGHV1-8*, *IGHV3-64D*, *IGHV3-9* and *IGHV5-10-1* usage, respectively (Fig. 6c). In both of these two examples, the GWAS SNVs and guQTLs were in strong LD with SVs spanning these respective sets of candidate genes (r = .51 and r=.98) suggesting that the observed effects could at least in part be SV-mediated.

**Figure 6.**
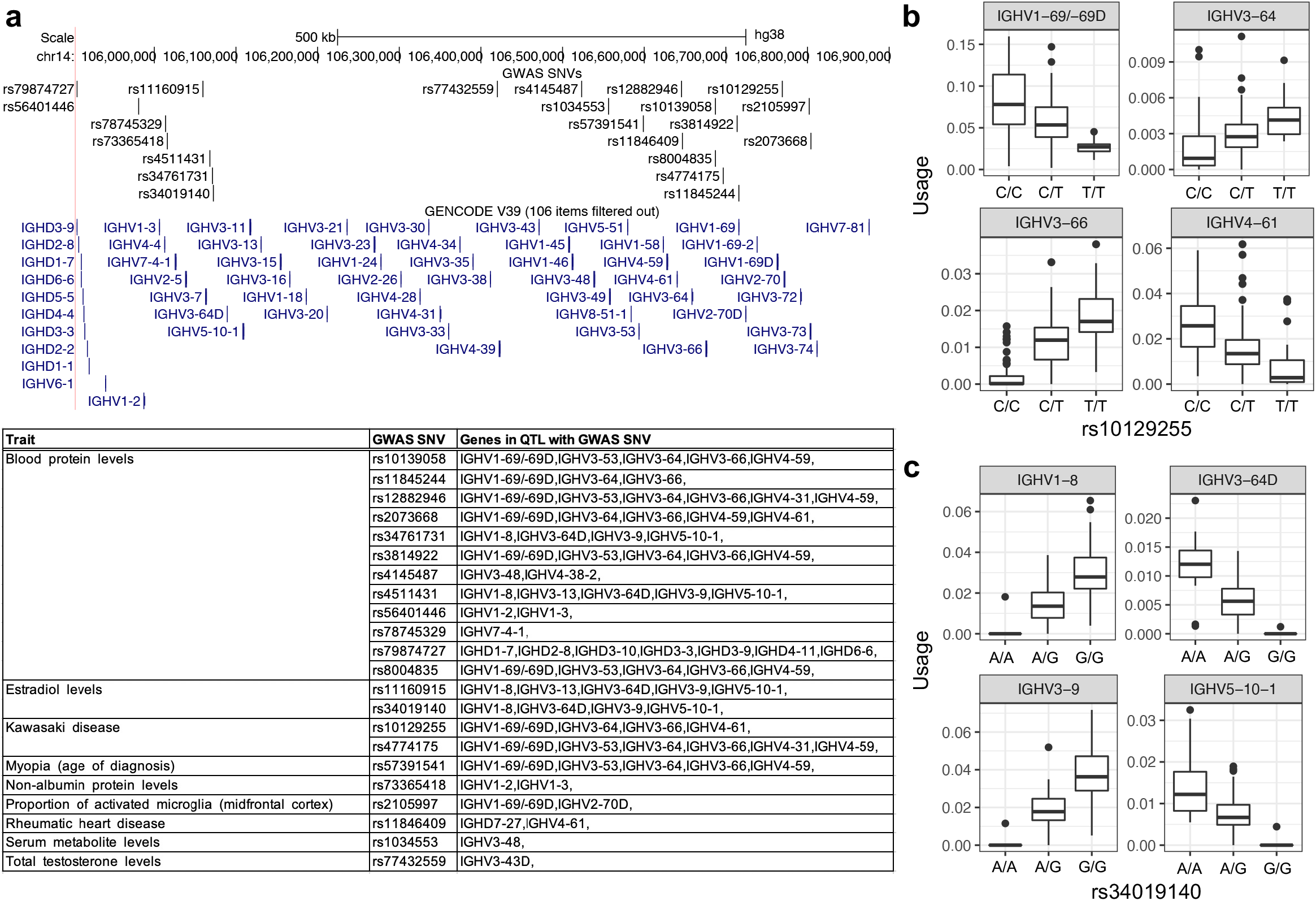
SNVs associated with diseases and traits are also associated with gene usage variation. **(a)** Map of IGH (GRCh38) showing the positions of SNVs identified by genome-wide association studies (GWAS); positions of F/ORF genes are also provided. For each GWAS SNV found to overlap a guQTL (IgM and IgG) from our dataset, the table provides information on the trait, SNV identifier, and genes for which usage was associated with the GWAS/guQTL SNV. **(b,c)** Box plots showing gene usage variation for all genes associated with two example GWAS SNVs for **(b)** Kawasaki disease and **(c)** estradiol levels.

### Repertoire-wide gene usage profiles are more highly correlated in individuals carrying shared IGH genotypes

Previous studies in monozygotic twins have shown that gene usage frequencies in genetically identical individuals are more highly correlated than in unrelated individuals^20, 21^. We reasoned that such effects could also be observed at the population level by assessing correlations in individuals sharing greater versus fewer IGH guQTL SNVs. To assess this, we used allele sharing distance^69, 70^ (ASD) to group individuals with similar genotypes across IGH and compare the IgM gene usage correlation between groups. Two ASD-based groupings were performed using either (1) the lead guQTL per gene (Fig. 7a), or (2) all guQTLs (Fig. 7b). We tested the latter case as we noted above that multiple variants could influence a single gene, and it has been shown that accounting for a greater number of common variants associated with a given phenotype can explain more variation in that phenotype^71^. Repertoire-wide gene usage correlations between samples were calculated using the Pearson’s Correlation coefficient. Using only the lead guQTL variants for each gene, individuals with the most overlapping guQTL genotypes (low ASD) had a higher mean IgM gene usage correlation than those in the group with the highest ASD scores (0.958 vs. 0.943; KS test *P value* < 3.8e-15). The same pattern was observed when using all significant variants (0.956 vs. 0.943; KS test *P value* = 0.008). These results indicated that genetic background makes a significant contribution to the overall gene usage composition of the repertoire, and expand on previous observations made in twin studies^20, 21^, by demonstrating that heritable components of the heavy chain repertoire can be directly linked to germline variants in the IGH locus.

**Figure 7.**
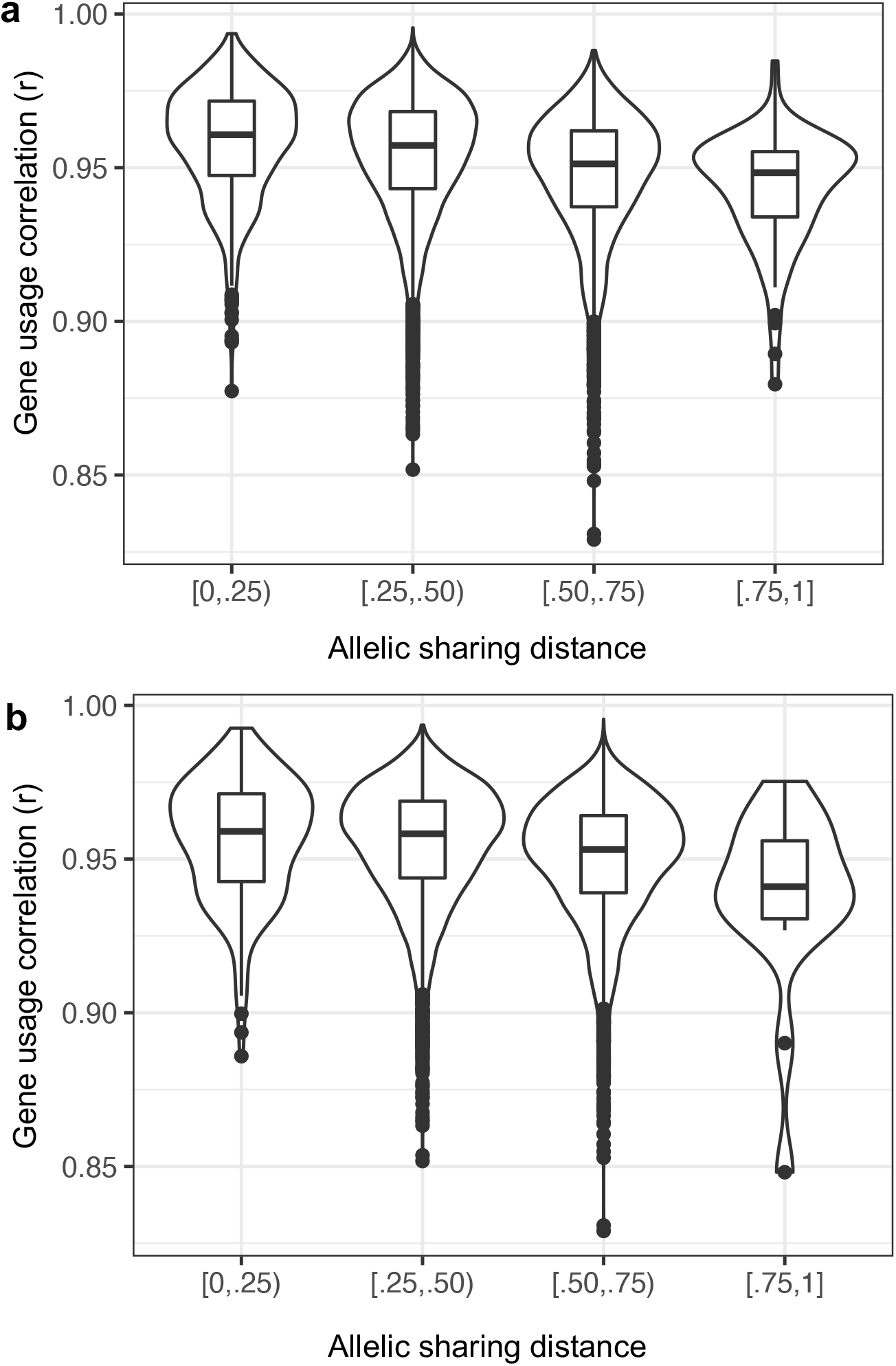
Individuals sharing a greater number of guQTL genotypes have more correlated repertoire-wide IgM gene usage profiles. **(a,b)** Pairwise intra-individual correlations (Pearson) of IgM usage for all genes, as well as allele sharing distance (ASD) for IGH SNV genotypes (lead guQTLs; all guQTLs) were calculated across individuals in the cohort. Violin plots show pairwise intra-individual repertoire-wide IgM gene usage correlations partitioned by ASD, calculated using either only lead guQTLs for all genes (**a**), or all guQTLs (**b**) for all genes (Bonferroni corrected).

## Discussion

In this study, we show conclusively that IGH genetic polymorphisms influence the composition of the Ab repertoire through impacts on gene usage frequencies. Resolution of complex IGH genetic variants using long-read sequencing identified associations between these variants and gene usage within the IgM and antigen-stimulated (IgG) repertoire. Variants were found to affect the Ab repertoire via (1) SVs that alter IGH gene copy number, including deletions that completely remove genes from the repertoire, as well as through (2) SNVs and indels, including those overlapping regulatory elements and transcription factor binding sites linked to V(D)J recombination. The strength of these associations was substantial, in some cases explaining >70% of variance in usage of particular genes. Building on past observations from twin studies^20, 21^, we found that repertoire-wide gene usage patterns were more similar in individuals sharing a greater number of genotypes across IGH. Together, these findings (1) advance our basic understanding of repertoire development, illuminating regions of IGH involved in gene regulation, and (2) more broadly represent a paradigm shift towards a model in which the Ab repertoire is formed by both deterministic and stochastic processes. This shift has critical implications for delineating the function of Abs in disease, with great potential to inform the design and administration of therapeutics and vaccines.

Resolving IG germline variants has historically been impeded by technical challenges resulting from the complex and highly polymorphic nature of the IG loci. Specifically, high-throughput approaches, including microarray and short-read sequencing are not able to fully and accurately resolve IG germline variation^28, 72^. Long-read sequencing has proven invaluable for resolving complex genomic regions, resulting in drastic improvements in variant detection^29, 73^. However, whole genome sequencing of large cohorts with long-read sequencing remains costly, laborious and prohibitive in many cases. Our alternative approach, using a targeted long-read protocol to selectively sequence the IGH locus in a cost-effective, multiplexed fashion, allowed us to characterize a broad spectrum of genetic variants in IGH for 154 donors, providing the largest long-read resolved collection of SVs, SNVs, and indels for this locus to date.

SVs are a hallmark of the IGH locus^25–27, 46, 74^, which was clearly supported by our analysis. We breakpoint resolved 28 SV haplotypes/alleles within 8 different SV loci spanning 542 Kbp of IGH; this included 13 novel SV alleles, and collectively resulted in copy number changes in 6 IGHD genes and 31 IGHV genes, representing 22% and 53% of all IGHD and IGHV genes in IGH, respectively. Critically, our ability to resolve SVs allowed us to more comprehensively detect and genotype SNVs and indels. In total, we identified 20,510 unique SNVs and 966 indels, 7980 and 223 of which were common. A significant fraction of these overlapped SVs (n=3,406), which we accurately genotyped as hemizygous. The increased performance of our approach was demonstrated through a comparison of our callsets to dbSNP, which revealed that the majority of common SNVs (63%; n=5057) detected were labeled as rare in frequency, lacked allele frequency data, or were completely missing from dbSNP altogether. Additional novelty was discovered through the annotation of IGH genes, revealing 135 undocumated alleles not currently curated in the germline gene database IMGT^75^. Together, these data hinted at the extent of variation that we have yet to describe in this complex locus, and bolster previous concerns that past genetic studies have overlooked IGH variants^28, 31, 76^. A major outcome of this study is that these data can start to be used to augment existing resources and databases that aim to provide improved reference data for the IG loci^30, 77^.

In addition to increasing our knowledge of IGH diversity, our ability to more fully resolve polymorphisms facilitated the identification of IGH germline variants that impact Ab repertoire diversity at a level that was previously not possible. Identifying associations between genetic variants and gene expression is a key step in determining the functional roles of germline variation in disease and clinical phenotypes, as well as resolving the molecular mechanisms that underlie gene regulation^48, 78^. By combining genetic variants with gene usage information across IGHV, D and J genes derived from AIRR-seq data, we performed the first gene usage QTL analysis, assessing associations between 7,297 common variants and 81 genes to identify polymorphisms explaining gene usage in the expressed IgM and IgG repertoire. These analyses revealed that half (52%) of common variants were associated with gene usage variation, impacting 59 (73%) genes in the IgM repertoire, with similar results in the IgG repertoire, indicating that patterns in IgG are likely highly influenced by the gene usage composition initially established in IgM, as noted previously^20, 21^. Furthermore, for 10 of the 59 genes identified in our analysis (9 of which were within SVs), the most significant variant explained more than half of the gene usage variation (*R*^2^ > .5). A conditional analysis further found that for 14 out of the 59 guQTL-associated genes in IgM, additional variance in gene usage could be explained by secondary polymorphisms, indicating that for at least a subset of IGH genes, interactions and additive effects across multiple variants will ultimately need to be resolved. These collective effects of polymorphisms across the repertoire as a whole were clear when we compared repertoires between individuals based on genetic similarity. As expected^20, 21^, we found that usage patterns were more highly correlated in individuals sharing IGH genotypes. This indicated that overlapping signatures in the repertoires of different individuals may be possible to identify and characterize with greater resolution at the population level by simply taking into account IGH genetic data^22^.

The guQTLs discovered provide the first insights into the potential functional mechanisms underlying the development of the Ab repertoire in humans. First, the association between SVs and gene usage variation offer a straightforward model for how germline variants impact the repertoire. Specifically, our results indicated that SVs change the copy number of genes, directly modifying their usage frequency in an additive fashion, likely by influencing the probability that the SV-associated genes are selected by V(D)J recombination based on the number of chromosomes on which they are present. This pattern was observed for the majority of genes associated with SVs in our dataset, and has been noted previously^40, 42^. Interestingly, there were also genes for which usage was impacted by neighboring SVs, even though the copy number of these genes was not directly altered, suggesting more complex mechanisms^42^. Beyond the effects of SVs, we found a significant number of SNVs associated with gene usage, all of which were in intergenic regions; again, this highlights the importance of our approach for capturing all IGH variant types, beyond just coding polymorphisms. Network analysis connecting genes with overlapping guQTL variants identified sets of genes whose usage patterns were coordinated; in many cases these genes were co-localized to specific regions of IGH, spanning 10’s to 100’s of Kbp. As with patterns observed for SVs, these signatures were illustrative of more complex regulatory mechanisms in the IGH locus. These regional effects appear consistent with studies of V(D)J recombination in model organisms. For example, the mouse IG loci partition into distinct regions, marked by specific regulatory marks, including TFBS and histone modification signatures, many of which, alongside RSS variation, have been associated with intra-gene V(D)J recombination frequency differences^32, 79, 80^. The mouse IG loci are also characterized by 3-dimensional structure, TADs and sub-TADs, associated with complex interactions between gene promoters and enhancers that coordinate V(D)J recombination in pre-B cells^35, 52, 81–83^. In contrast to mouse, functional genomic elements dictating V(D)J recombation in the human IGH locus have not been characterized in depth; nonetheless, our intersection of guQTLs with publicly available annotation sets revealed enrichments in cis-regulatory elements and TFBS involved in V(D)J recombination in animal models. This included CTCF and EED TFBS, as well as IGH regions marked by H3K4me3^53, 60–62^. While fine mapping and functional validation of guQTLs is needed, this result was reaffirming given that gene usage in the IgM repertoire is a proximal measurement of V(D)J recombination, providing initial evidence that the variants we identified likely influence the frequency at which IGH genes are selected during V(D)J recombination.

Ultimately, an improved understanding of Ab repertoire diversity and function will be critical to resolving the role of B cells in disease. This study provides support for the idea that leveraging IG genetic data can better delineate Ab response dynamics in a variety of contexts. For one, there is growing interest in developing predictive models for V(D)J recombination and repertoire diversity^84, 85^, and applying Ab repertoire profiling as a diagnostic tool for disease and clinical phenotypes of high public health relevance^86, 87^. However, current models do not explicitly account for genetic factors, and the effects of this on model performance are not known^84, 85^. Our results indicate that future work in this area should explore ways to integrate genetic data; this will likely be critical for better understanding commonalities and differences in repertoire signatures, not only for gene usage patterns, but also in identifying additional features (e.g., public clonotypes^1,2^), overall leading to improved metrics for immune response monitoring and prediction modeling.

Here, we demonstrate that our data already provide an opportunity to more fully explore the potential roles of IGH polymorphism in Ab-mediated diseases. First, the direct overlap of GWAS SNVs and guQTLs indicate the potential for effects of GWAS variants to be mediated through genetic effects on Ab gene usage. This parallels approaches employed for eQTLs and GWAS variants elsewhere in the genome to nominate genes/pathways underlying human phenotypes^48, 88–90^. As additional disease genetic associations are made in IGH, our dataset will continue to be useful for making such first-line connections, and drive the generation of novel hypotheses that can be explored experimentally. Second, our results can directly inform our understanding of vaccine responsiveness, particularly as this pertains to efforts centered around the elicitation of targeted antibodies. Notably, our analysis revealed that IGHV coding variation was in many cases linked to guQTLs, indicating that usage patterns can coincide with amino acid differences that are important for Ab-antigen interactions. This is consistent with previous reports^23, 41, 91^, including examples related to precursor germline alleles critical for broadly neutralizing Abs in various infectious diseases. For example, it has been shown that IGHV1-2 germline alleles associated with HIV VRC01 Abs, which are a current focus of germline targeting immunogens, have variable usage frequencies in the IgM repertoire and associate with variable immunogen-specific B cell frequencies^23^. Another clear example is a germline variant that encodes a critical phenylalanine within the CDR2 of IGHV1-69-derived broadly neutralizing Abs against the influenza hemagglutinin stem^41, 64^. This variant has not only been shown to facilitate antigen binding, but also (mirroring patterns observed for VRC01 alleles) is associated with variable usage patterns in the IgM and IgG repertoire^41^. Interestingly, in both of these examples, allelic variants vary considerably between human populations^23, 41^, indicating that both population-level diversity and the role of germline variants in shaping the baseline B cell repertoire will need to be considered in interpreting vaccine response data^22^.

While the dataset we have analyzed here represents the most comprehensive survey to date, it is likely that increasing the sample size will uncover additional genetic contributions to gene usage. For example, by lowering our *P* value threshold by only a factor of 10, the fraction of IGH genes with usage associated to at least one genetic variant increased from 73% to 91% (74/81). This further bolsters our finding that a large fraction of variation in repertoire gene usage between individuals will likely be explained by variants in IGH. Rarer and complex IGH variants will need to be better accounted for in future work, specifically those excluded from our analysis due to low frequency and genotyping coverage. For example, SV alleles within the highly complex and polymorphic IGHV3-30 region will require sequencing and haplotyping in larger cohorts to better resolve the effects of variation for those genes, which have suspected roles in disease^92, 93^. In addition, it will be important for future work to also consider integrating analyses of the IG light chain loci. Light chain genes contribute to Ab folding and Ab-antigen interactions^94–96^, and it is plausible that both trans-effects and interactions between heavy and light chain variants could influence gene usage. The development of models that incorporate both genetic variation and features specific to both chains (e.g., binding and stabilization), would more fully delineate the total genetic contribution to variation in the Ab repertoire. In addition, as cohorts increase in size, additional insight will come from the consideration of other variables such as genetic ancestry, positive/negative selection, age, B cell subset and tissue^97–99^. Finally, the models utilized here could be extended to assess the contribution of IGH polymorphisms to other repertoire signatures, including N/P addition and CDR3 features, which also are influenced by heritable factors^20, 21, 38, 85^.

Collectively, our analyses provide the first comprehensive picture of IGH polymorphism and Ab repertoire variation. These findings have the potential to reshape the way we conduct, analyze and interpret AIRR-seq data, and use these data to profile the Ab response in disease. As noted previously, the results provided here further illuminate the need for improving efforts to more fully explore the extent of IGH polymorphism in the human population, as a means to resolve the role of germline variation in Ab function and disease.

## Materials and Methods

### Long-read library preparation and sequencing

Genomic DNA was extracted from peripheral blood mononuclear cells (PBMC) or polymorphonuclear neutrophil (PMN) procured from Stanford University, Harvard University or STEMCELL Technologies (Vancouver, Canada). Genomic DNA was processed using our published targeted long-read sequencing protocol^28^. High molecular weight DNA (0.5-2 ug) was sheared using g-tubes (Covaris) and size selected using the 0.75% DF 3-10 Kbp Marker S1-Improved Recovery cassette definition on the Blue Pippin (Sage Science); library size ranges provided in Supplementary Fig. 1. The DNA was End Repaired and A-tailed using the standard KAPA library protocol (Roche). Barcodes were added to samples sequenced in multiplex pools and universal primers were ligated to all samples. PCR amplification was performed for 8-9 cycles using high-fidelity polymerase (LA-Taq or PrimeSTAR GXL, Takara) at an annealing temperature of 60℃. Small fragments and excess reagents were removed using 0.7X AMPure PB beads (Pacific Biosciences). Libraries were hybridized to IGH-specific oligonucleotide probes (Roche; see reference^28^) and recovered using streptavidin beads (Life Technologies) prior to another round of PCR amplification for 16-18 cycles using either LA-Taq or PrimeSTAR GXL (Takara) at an annealing temperature of 60℃.

Enriched IGH libraries were prepared for sequencing using the SMRTbell Express Template Preparation Kit 2.0 (Pacific Biosciences). DNA was treated with Damage Repair and End Repair mix to repair nicked DNA, followed by the addition of an A-tail and overhang ligation with SMRTbell adapters. These libraries were treated with a nuclease cocktail to remove unligated input material and cleaned with 0.45X AMPure PB beads (Pacific Biosciences). The resulting libraries were prepared for sequencing according to the manufacturer’s protocol and sequenced as single libraries per SMRTcell with P6/C4 chemistry and 6h movies on the RSII system, or as multiplexed libraries sequenced on the Sequel (3.0 chemistry; 20h movies) or Sequel II/IIe system (2.0 chemistry; 30h movies).

Generated targeted capture libraries had an average insert length of 6 Kbp, and were sequenced using the Pacific Bioscience (PacBio) RSII (n=40), Sequel (n=40) or Sequel IIe (n=74) systems (Table 1). This strategy confers two main advantages: (1) the sequencing polymerase passes over amplicons multiple times, allowing for the generation of highly accurate (high-fidelity, HiFi) reads (Supplementary Figure 1a,b); and (2), for Sequel/IIe libraries, multiple samples are barcoded and sequenced in a single sequencing run. Critically, the high HiFi read quality overcomes historical concerns of high error rates in long-read sequencing data (Table 1), and error-correction steps performed during the assembly process increases the read base-level accuracy^100, 101^. Previously, we have shown that assemblies produced from the older RSII platform have high base-level accuracy^28^.

For a single sample, we prepared libraries for adaptive nanopore sequencing using the Ligation Sequencing Kit (Oxford Nanopore Technologies, ONT) and the NEBNext Companion Module for ONT Ligation Sequencing (New England Biolabs). 3 μg gDNA was used as input for these libraries. Entire purified libraries (5-50 fmol, per manufacturer’s recommendation) were loaded onto R9.4.1 flow cells on the MinION Mk1C instrument (ONT). The experimental run was set up with no multiplexing, turning on enrich.fast5, and using human nanopore enrichment. Additionally, fast (or high accuracy) base calling was employed for a 72-hour run. In addition to IGH, multiple genomic loci were targeted for sequencing in order to provide the minimum number of bases (17 Mb) required for adaptive sequencing. The IGH sequence targeted was from the custom reference used in this study (below).

### IgG and IgM antibody repertoire sequencing

For newly generated expressed Ab repertoire sequencing datasets, total RNA was extracted from PBMCs using the RNeasy Mini kit (Qiagen). For each sample, IgG and IgM 5’RACE AIRR-seq libraries were generated using the SMARTer Human BCR Profiling Kit (Takara Bio), following the manufacturer’s instructions. Individually indexed IgG and IgM libraries were assessed using the Agilent 2100 Bioanalyzer High Sensitivity DNA Assay Kit (Agilent) and the Qubit 3.0 Fluorometer dsDNA High Sensitivity Assay Kit (Life Technologies). Libraries were pooled to 10 nM and sequenced on the Illumina MiSeq platform using the 300bp paired-end reads with the 600-cycle MiSeq Reagent Kit v3 (Illumina). Additional datasets were downloaded from SRA for Nielsen et al^18^ and Ke et al (unpublished).

### Custom linear IGH reference

A custom linear reference for IGH was used that includes previously resolved insertion sequences^25^ absent in GRCh38. This reference was previously used and vetted to generate high confidence variant call sets^28^. The reference was built off of GRCh38 (chr14:105860500-107043718). Partial sequences from GRCh38 were removed and additional insertion sequences were added from previously characterized structural variants^25^. Specifically, sequence between chr14:106254581-106276923 (GRCh38) was swapped for a 10.8 Kbp duplication containing the *IGHV3-23D* gene from fosmids ABC9-43993300H10 and ABC9-43849600N9. Sequences between chr14:106317171-106363211 (GRCh38) and chr14:106403456-106424795 (GRCh38) was swapped for a 77.6 Kbp duplication haplotype containing IGHV genes *IGHV3-30, IGHV4-30-2*, *IGHV3-30-3*, *IGHV4-30-4*, *IGHV3-30-5*, *IGHV4-31* and *IGHV3-33* from fosmid clones ABC11-47150400I4, ABC11-47354200D2 and ABC11-49598600E10, and a 75.8 Kbp insertion containing IGHV genes *IGHV3-38 IGHV4-38-2*, *IGHV3-43D, IGHV3-38-3*, *IGHV1-38-4* and *IGHV4-39* from fosmid clones ABC10-44084700I10, ABC10-44145400L1 and WI2-1707G1, respectively. A 37.7Kbp complex SV with *IGHV3-9* and *IGHV1-8* genes derived from GRCh37 (chr14:106531320-106569343) was appended to the end of the reference separated by 5 Kbp of gap sequence (“N”). This reference sequence is available on github (https://github.com/oscarlr/IGenotyper).

### IGH locus assembly and variant detection

All targeted long-read datasets were processed using IGenotyper with default parameters^28^. IGenotyper uses BLASR^102^, WhatsHap^103^, MsPAC^104^ and Canu^100^ to align reads, call and phase SNVs, phase reads and assemble phase reads, respectively. Using the assemblies, IGenotyper uses the MsPAC multiple sequencing alignment and Hidden Markov model module to identify SNVs, indels and SVs. SVs not resolved were genotyped using HiFi read coverage and soft-clipped sequences in the assembly and in HiFi reads, and manually resolved using BLAST and custom python scripts. All SV genotypes were visually inspected using Integrated Genome Viewer (IGV) screenshots generated from an IGV batch script.

### Characterizing novel alleles and expanding the IGH allele database

Novel alleles for IGHV, IGHD and IGHJ genes supported by 10 HiFi reads (exact matches) or found in 2 or more individuals were extracted from the assemblies of each sample. Novel alleles were defined as those not found in the IMGT database (release 202130-2). Allele sequences that aligned to IMGT alleles with 100% identity were also characterized as novel, if the putative novel allele was annotated from a gene in the assembly that was different from the gene assignment in the IMGT database. The non-redundant set of novel alleles was appended to the IMGT database for IgM/IgG repertoire sequencing analyses conducted in this study. A BLAST database was created using makeblastdb version 2.11.0+. Gapped sequences for the novel alleles were generated using the IMGT/V-QUEST server^105^.

### Processing AIRR-sequencing data

Paired-end sequences (“R1” and “R2”) were processed using the pRESTO toolkit^106^. All R1 and R2 reads were trimmed to Q=20, and reads <125 bp were excluded using the functions “FilterSeq.py trimqual” and “FilterSeq length”, respectively. Constant region (IgM and IgG) primers were identified with an error rate of 0.2 and corresponding isotypes were recorded in the fastq headers using “MaskPrimers align”.

For sequencing datasets without unique molecular identifiers (UMIs), R1 and R2 reads were assembled using “AssemblePairs align”, and resulting merged sequences <400 bp were removed using “FilterSeq length”. Identical sequences were collapsed, and read duplicate counts (“Dupcounts”) were recorded. For sequencing datasets with UMIs, the 12 base UMI, located directly after the constant region primer, was extracted using “MaskPrimers extract”. Sequences assigned to identical UMIs were grouped and aligned using “ClusterSets” and “AlignSets muscle”, and then consensus sequences were generated for each unique UMI set using “BuildConsensus”. Identical sequences with different UMIs were collapsed, and read duplicate counts (“Dupcounts”) were recorded. Collapsed consensus sequences represented by <2 reads were discarded.

Processed AIRR-seq fastq files were split by isotype using the “SplitSeq.py group” function from Immcantation^106^. Samples with <100 reads per isotype were removed. Following the application of this filter, the mean number of merged consensus sequences per repertoire ranged from 465 to 109250 (mean=26036), with lengths ranging from 318 to 510 bp. Fastq files were aligned to the expanded database, including IMGT and novel alleles identified in our cohort, using “AssignGenes.py igblast” to generate Change-O^107, 108^ files. Productive reads were specifically selected using the “ParseDb.py split” command. Assignments to genes found to be deleted from both chromosomes in genomic datasets for a given sample were removed from the Change-O. Reads assigned to multiple alleles were re-assigned to a single allele if and only if the genomic data revealed that only one of the alleles was present. Clones were detected using the modified Change-Os with the ‘shazam distToNearest’ command and ‘model=’ham’, normalize=’len’’ parameters, ‘shazam findThreshold (parameters: method=’gmm’, model=’gamma-gammà), and ‘DefineClones.py (parameters: –act set –model ham –norm len –mode allele)’ commands. IgM and IgG repertoires with fewer than 200 clones identified were excluded from downstream analysis.

### Calculating gene usage among defined clones

A *m x n* clone count matrix *C* was created, where *m* are the genes and *n* are the samples. Due to the sequence similarity, duplicated genes were summed into a single entity. The counts of the following genes combined:

1. IGHV3-23 and IGHV3-23D
2. IGHV3-30, IGHV3-30-3, IGHV3-30-5 and IGHV3-33
3. IGHV1-69 and IGHV1-69D
4. IGHD4-4 and IGHD4-11

*C* was batch corrected (3 batches) using ComBat-seq^109^ to produce an adjusted count matrix *C*′ to account for differences between the three AIRR-seq datasets used. The fractions of clones per gene or gene set (*m*) was calculated from *C*′ across each sample (*n*).

The following set of F/ORF genes were removed or not analyzed:

1. *IGHD5-5*: In all cases where *IGHD5-5* was identified through IgBLAST, the AIRR-seq reads were assigned to *IGHD5-5*01* and *IGHD5-18*01*, or *IGHD5-5*01*, *IGH5-18*01* and additional alleles. The genes *IGHD5-5* and *IGHD5-18* were not combined because there were AIRR-seq reads aligned solely to *IGHD5-18*.
2. *IGHV3-16*: No AIRR-seq reads aligned to *IGHV3-16*.

### Selecting common variants for gene usage QTL analysis

SNVs with a HWE value less than 0.000001 were filtered using bcftools^110^. SNVs found in less than 5 individuals were removed if they did not have HiFi read support. TheSNVs passing these stringent quality control thresholds were used to impute missing genotypes using Beagle^111^ (v228Jun21.220). The resulting SNVs were again filtered if they contained a HWE value less 0.000001. Common SNVs were selected if they were genotyped in at least 40 individuals and had a MAF equal to or greater than 0.05. The same criteria were applied to SNVs selected for conditional analysis.

Indels and SVs, excluding large SVs (> 9 Kbp), were split into two categories based on whether they overlapped tandem repeat regions. Tandem repeat regions on the custom reference were determined using Tandem Repeats Finder^112^ with parameters (match = 2, mismatch = 7, delta = 7, PM = 80, PI =10, Minscore = 10, MaxPeriod=2000). Events overlapping tandem repeats were genotyped again in all the samples using the dynamic programming algorithm from PacMonSTR^113^. Events were merged using a custom python script (https://github.com/oscarlr-TRs/PacMonSTR-merge). Tandem repeat events with an alignment score between the motif and the copies in the assemblies lower than .9 were removed. Tandem repeat alleles were defined by a difference of a single motif copy. Tandem repeat events with an allele occurring at a frequency greater than 0.05 was considered common. An expansion or contraction greater than 50 bps relative to the reference was considered a tandem repeat SV. Indels and SVs from IGenotyper outside of tandem repeats across all samples were merged. Manual inspection showed high concordance between event sizes and sequence content. In cases where a discordance was observed between event sizes, the max size was selected. Samples were genotyped as homozygous reference for indels and SVs if no event was detected and both haplotypes were assembled over the event. Indels and SVs with a MAF greater than 0.05 were selected.

All SVs were genotyped using IGenotyper and manually inspected using IGV. SVs with a MAF less than 0.05 were not included in the QTL analysis (Supplementary Table 2).

### Gene usage QTL analysis

SNVs, complex SVs and mSVs were associated with usage using ANOVA and linear regression. All other variant types, indels, non complex SVs and large SVs (excluding mSVs) were associated with usage using linear regression. Both models included age and AIRR-seq sequencing platform as covariate (n=3). A linear regression was used to extract additional metrics (e.g. beta, R^2^). Associations were corrected for multiple hypothesis testing using Bonferroni correction on a per-gene level. Variants with an LD of 1 were treated as a single variant during correction. Conditional analysis was performed in the same manner using all variant types with the same filters applied to the initial call sets.

### Network analysis of variants associated with multiple genes

Variant and gene pairs for variants significantly associated with more than 1 gene in the IgM repertoire were selected. A graph using the networkx python library (networkx.org) was created with genes as nodes and edges connecting genes/nodes if the same variant was associated with both genes. An edge weight was given for each time nodes were connected. The graph was pruned such that the edge weights were greater than 2. Cliques were identified using the find_cliques function.

### Regulatory analysis

ENCODE cCREs were downloaded from the UCSC Genome Browser under group “Regulation”, track “ENCODE cCREs” and table “encode CccreCombined”. ENCODE transcription factor binding site data were also downloaded from the UCSC Genome Browser under group “Regulation”, track “TF Clusters” and table “encRegTfbsClustered”. SNVs associated with gene usage were overlapped with both tracks and an enrichment in both tracks over all SNVs overlapping each track was calculated using a one-sided Fisher Exact Test.

### GWAS analysis

Variants identified by GWAS with an association *P value* lower than 4e-6 were downloaded from the NHGRI-EBI GWAS catalog (https://www.ebi.ac.uk/gwas/api/search/downloads/full). Significant variants from this study were intersected with GWAS variants.

## Author contributions

OLR, MLS, WAM, and CTW conceived and planned the study. OLR, YS and DT performed computational experiments. CAS, KS, WSG, JT HK and KJLJ performed wet lab experiments. WAM, MLS and CTW supervised the study. HK, KJLJ, SB, WAM and CTW provided samples and data. All authors read and approved the final manuscript.

## Data Availability

The datasets generated during and/or analysed during the current study are available from the corresponding author on reasonable request.

