## Supplementary Figures for "Genetic variation in the immunoglobulin heavy chain locus shapes the human antibody repertoire"

### Supplement

### Results

#### Detailed descriptions of structural variants

The two largest SV alleles occurred within a single mSV and were 259 Kbp and 284 Kbp long, resulting in deletions of 14 and 16 IGHV genes, respectively (Fig. 1b). These deletions were observed only in White individuals (n=7). This observation is likely explained by the fact that one of the segmental duplication blocks that mediates these deletions occurs on a complex SV allele with genes *IGHV3-64D* and *IGHV5-10-1*, which is found at higher frequencies in European populations[1]. These large deletions have been partially resolved from AIRR-seq data[2], giving further support to their authenticity.

The region surrounding *IGHV3-30* and *IGHV4-28* and related genes (*IGHV4-30-2, IGHV3-30-3, IGHV4-30-4, IGHV3-30-5, IGHV4-31* and *IGHV3-33*) has been identified previously as a SV hotspot[3]. In earlier studies, 4 SV alleles in this region were fully resolved[3,4]. The longest resolved SV allele spans ~100 Kbp and harbors 4 ~25 Kbp segmental duplications, consisting of repeating IGHV4 and IGHV3 gene cassettes. In this study, we observed 4 of the previously characterized SV alleles, as well as 8 novel SV alleles (Fig. 1c). Relative to the longest SV allele, the other 11 SV alleles contained deletions that varied by position and ranged in size from 23.9 to 74.2 Kbp.

The other SV hotspot identified was an mSV with 4 SV alleles spanning 136 Kbp and included the genes *IGHV4-38-2*, *IGHV3-43D, IGHV3-38-3, IGHV1-38-4, IGHV4-39* and *IGHV3-43* (Fig. 1d). The SV allele harboring all of these genes is present in our custom reference and was previously resolved[3]. In addition to this haplotype, we identified three deletions (two novel) and one insertion containing two newly discovered paralog genes with 100% sequence identity to *IGHV4-38-2*02* and *IGHV3-43D*03*. Self-alignment of the haplotype with the insertion to itself further identified that the ~52.2 Kbp insertion is a partial duplication of a previously resolved SV allele[3] (Supplementary Figure 4a,b). Additionally, we employed adaptive ("read-until") nanopore sequencing in combination with the targeted HiFi long-read sequencing derived assemblies to fully span this event (Supplementary Figure 4c,d).

In addition to the previously characterized SV allele including *IGHV3-23* and *IGHV3-23D*, we identified a duplication that contained three *IGHV3-23* gene copies (Fig. 1e). Out of the 6 individuals carrying this duplication, 5 were Asian. A higher *IGHV3-23* gene copy number in Asians was reported previously in an early restriction fragment length polymorphism study[5].

While many SVs have been characterized in the IGHV gene region, SVs within the IGHD gene region have usually been predicted using AIRR-seq data, with very limited evaluation by germline locus sequencing[6,7]. Critically, IGHD genes make up a large portion of the complementary determining region 3 (CDR3), the most somatically variable Ab region[8] and a critical determinant of antigen specificity[9]. In our cohort, we characterized a previously inferred deletion spanning 9.6 Kbp, deleting 6 (*IGHD2-8, IGHD1-7, IGHD6-6, IGHD5-5, IGHD4-4, and IGHD3-3*) out of the 26 (23%) IGHD genes (Fig. 1f). Interestingly, this deletion was common (allele frequency = 0.19), present in 23 out of 76 individuals for which genotyping was possible, and homozygous in 5 individuals. One of the homozygotes was also heterozygous for the largest 284 Kbp deletion in the IGHV gene region (Fig. 1b). Just between these two SVs, this individual carried a unique IGH haplotype with 6 deleted IGHD genes and 16 IGHV deleted genes. Taking into account the other SVs concurrently observed, an additional 13 IGHV genes were deleted, totaling 35 deleted IGHV and IGHD genes across both haplotypes in this individual (Fig. 1g).

#### Biased discovery of novel alleles in self-reported non-White individuals

The number of novel alleles identified across the cohort was not equally distributed among individuals. The majority of individuals (n=125; 81%) contained at least one novel allele, with 76 (61%) individuals having 1 to 3 novel alleles. Of the 8 individuals who had 10 or more novel alleles, 5 self-reported as Black or African American. Of the 35 individuals who had 5 or more novel alleles, 14, 7, 4, 2 and 6 were Black or African American, White, South Asian, East Asian and Hispanic or Latino, respectively. This corresponds to 70%, 8%, 20%, 18% and 32% of individuals from each respective subgroup. Furthermore, of the 25 novel alleles found in 5 or more individuals, 3 were found specifically in one subgroup. These novel alleles corresponded to genes *IGHV3-30-3*, *IGHV1-38-4* and *IGHV1-69D*, which are all found within SVs. Additionally, each of these novel alleles appeared at a high frequency, with the novel alleles for *IGHV3-30-3*, *IGHV1-38-4* and *IGHV1-69D* found in 8 Asian, and 7 and 5 Black or African American individuals, respectively.

### Supplementary figure legends

**Figure 1. PacBio sequencing and assembly statistics.**

**Figure 2. Number and length of merged reads for each AIRR-seq dataset after processing.**

**Figure 3. IGenotyper assembly and PacBio HiFi (capture) read coverage for two example SVs. (a)** IGV screenshot showing the assemblies and HiFi read profiles from two hemizygous individuals spanning the 9.6 Kbp deletion in the IGHD region. The positions of IGHD genes, including those deleted by the SV (IGHD2-8, IGHD1-7, IGHD5-5, and IGHD3-3) are provided. **(b)** HiFi coverage and assemblies for a sample homozygous for a mSV deletion allele including the genes *IGHV3-38-3*, IGHV1-38-4, IGHV4-39, and *IGHV3-43*. Green and blue HiFi reads and assemblies correspond to reads derived from the maternally or paternally haplotypes. Pink HiFi reads are reads that could not be assigned to either haplotype.

**Figure 4. Resolving a novel insertion with adaptive read-until Oxford Nanopore sequencing. (a)** A schematic showing the haplotype with the resolved ~52.2 Kbp insertion. The positions of the insertion (black bar) and IGHV genes are shown. Specifically, the insertion includes duplications of the genes IGHV4-38-2, and IGHV3-43D, which match known alleles for these genes in IMGT at 100% identity. **(b)** Dot plot of the self-alignment of the SV haplotype reveals that the region spans a large segmental duplication, including the insertion sequence, which represents a ~52 Kbp duplication block (red arrow), which occurs twice in this haplotype. **(c)** HiFi reads aligned to a shorter SV allele identified a read profile demonstrating a duplication. Green and blue HiFi reads and assemblies correspond to reads derived from the maternally or paternally haplotypes. Pink HiFi reads are reads that could not be assigned to either haplotype. This is usually because the reads correspond to a hemizygous deletion or are from a homozygous locus. Reads in the blue box correspond to a deletion haplotype shown in **(d)**. Reads derived from the alternate haplotype  show heterozygous SNVs , a typical signature of duplications. Using longer reads derived from adaptive read-until methods ("ONT reads" panel) in combination with the HiFi reads we were able to manually reconstruct the ~52 Kbp insertion. Arrows indicate the order in which ONT reads were combined to reconstruct the insertion haplotype shown in (A).

**Figure 5. Number of SNVs in different gene components across all IGHV genes.**

**Figure 6. Examples of polymorphic indels and small SVs. (a)** A polymorphic 86 bp variable number tandem repeat (VNTR) upstream of *IGHV3-20*. Across the cohort most individuals have 4 copies of the motif, however, some individuals have up to 9 copies. **(b)** A complex SV upstream of *IGHV1-3* is most likely derived from a degenerate tandem repeat. Three different SV alleles were identified with different motif sequences. The number of motif copies differed between SV alleles. An alignment of the consensus motif sequence to all the motifs in the SV alleles showed low sequence identity. **(c)** Polymorphic homopolymer expansion. The distribution of "T" copies, assemblies and HiFi reads are shown.

**Figure 7. Gene usage association statistics for the IgG repertoire.**

**Figure 8. Comparison of the IgM and IgG gene usage association results. (a)** The overlap of gene usage QTL genes between IgM and IgG. **(b)** The overlap of gene usage QTL variants between IgM and IgG. **(c)** Gene usage correlation between the IgM and IgG repertoire.

**Figure 9. Gene usage for genes in the largest deletion identified split by the genotype of the largest deletion.** Individuals with the deletion (genotype group “0”) had overall less usage than individuals without the deletion for genes in the deletion.

**Figure 10. Gene usage for individuals with different *IGHV3-23* copy number.**

**Figure 11. Gene usage for *IGHD3-10* is associated with distant IGHD gene region deletion even though it is outside of the IGHD gene deletion.**

**Figure 12. Network of all genes connected if a SNV is associated with both genes.**

**Figure 13. Cliques found in the network. (a)** Cliques containing mostly SV genes, **(b)** clique containing an equal amount of SV and non-SV genes and **(c)** cliques without mostly SV genes.

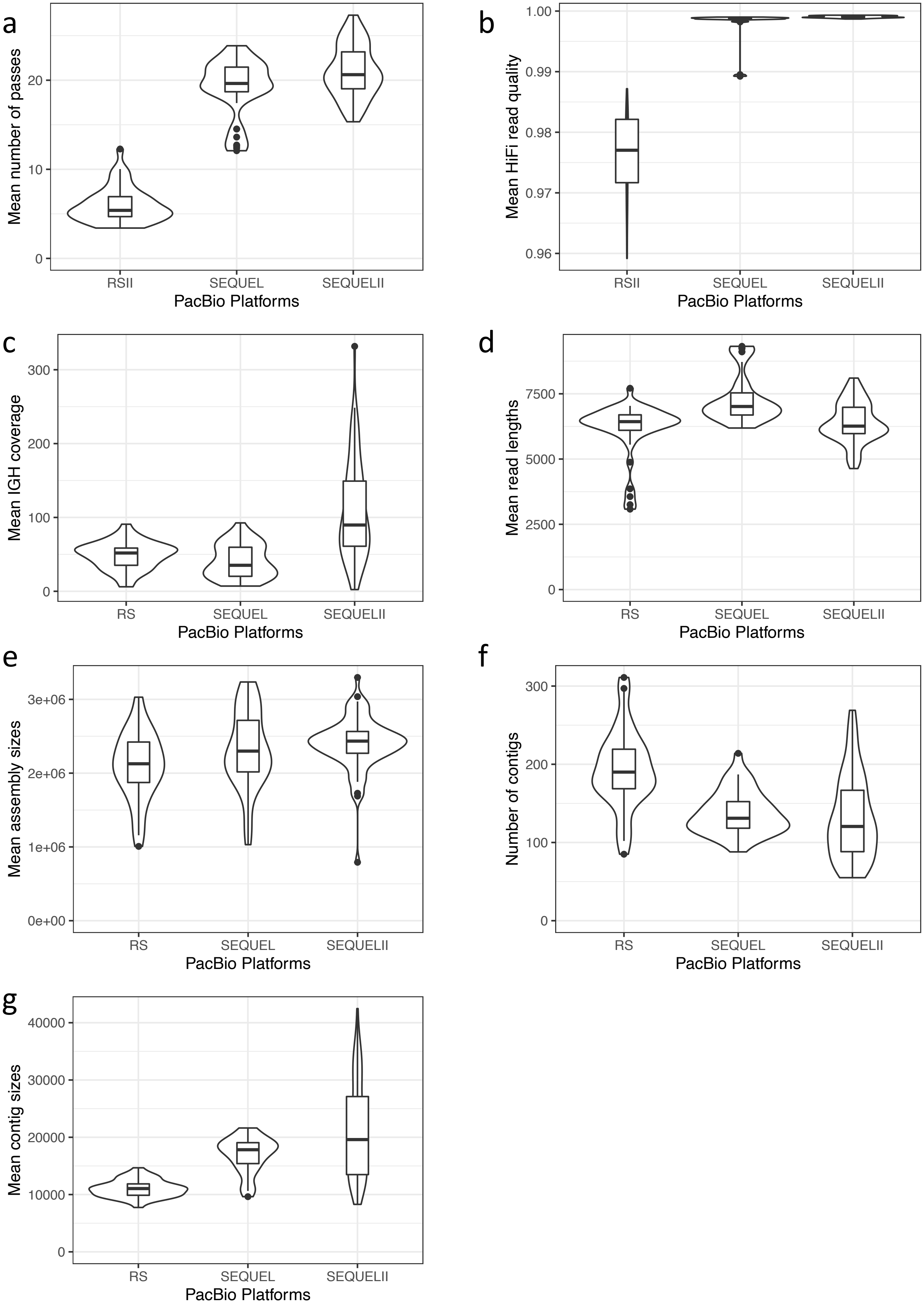
a
Mean number of passes
0
10
20
RSII
SEQUEL
SEQUELII
PacBio Platforms
b
Mean HiFi read quality
0.96
0.97
0.98
0.99
1.00
RSII
SEQUEL
SEQUELII
PacBio Platforms
c
Mean IGH coverage
0
100
200
300
RS
SEQUEL
SEQUELII
PacBio Platforms
d
Mean read lengths
0
2500
5000
7500
RS
SEQUEL
SEQUELII
PacBio Platforms
e
Mean assembly sizes
0e+00
1e+06
2e+06
3e+06
RS
SEQUEL
SEQUELII
PacBio Platforms
f
Number of contigs
0
100
200
300
RS
SEQUEL
SEQUELII
PacBio Platforms
g
Mean contig sizes
0
10000
20000
30000
40000
RS
SEQUEL
SEQUELII
PacBio Platforms

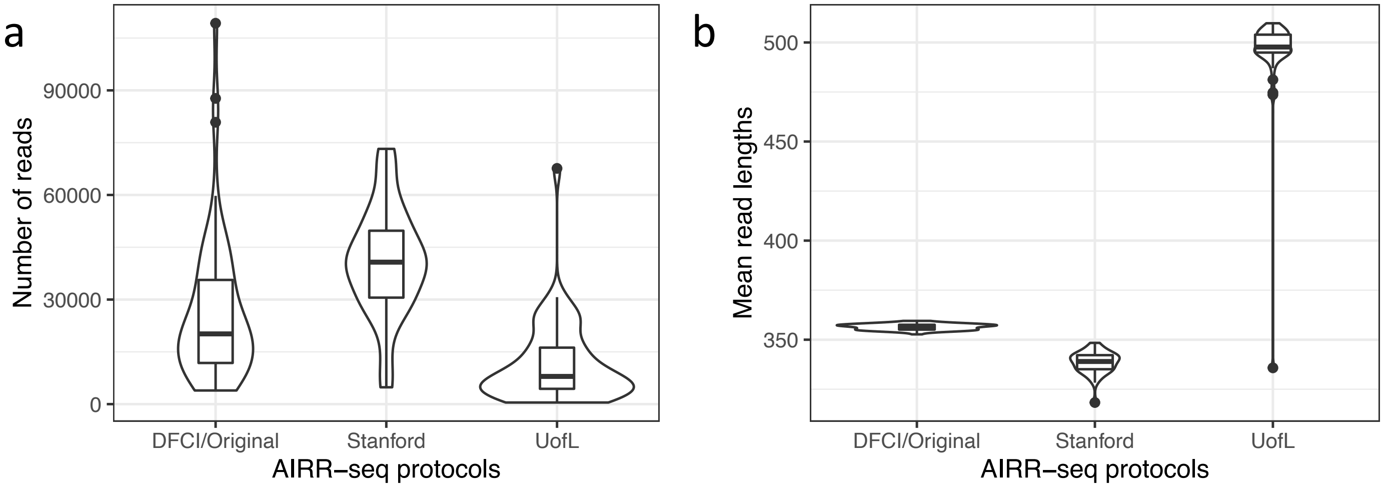
a
Number of reads
90000
60000
30000
0
DFCI/Original
Stanford
UofL
AIRR−seq protocols
b
Mean read lengths
500
450
400
350
DFCI/Original
Stanford
UofL
AIRR−seq protocols

a

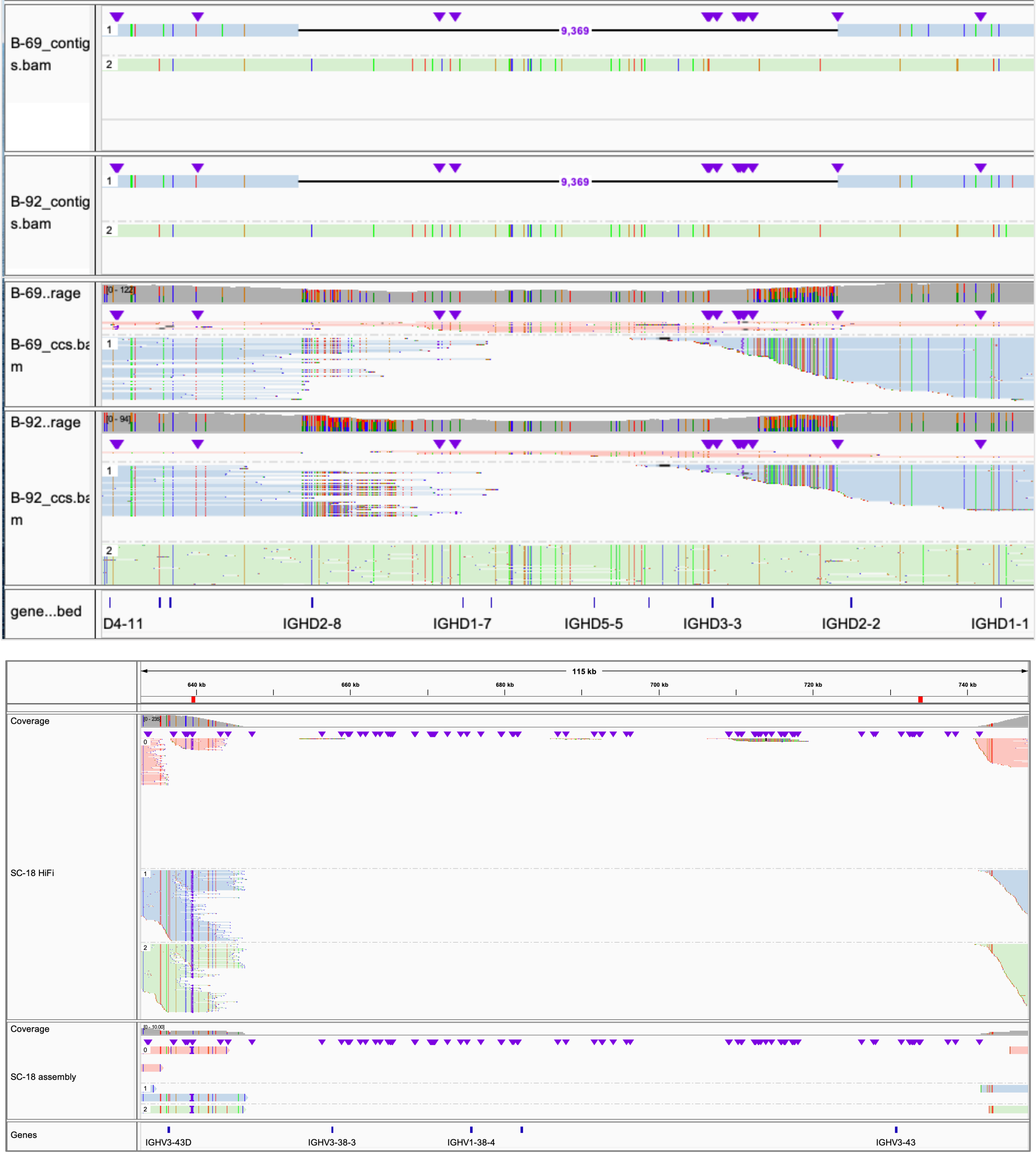

B-69_contigs.bam
9,369
B-92_contigs.bam
9,369
B-69..rage
[0 - 122]
B-69_ccs.bam
B-92..rage
[0 - 94]
B-92_ccs.bam
gene...bed
D4-11
IGHD2-8
IGHD1-7
IGHD5-5
IGHD3-3
IGHD2-2
IGHD1-1
115 kb
640 kb
660 kb
680 kb
700 kb
720 kb
740 kb
Coverage
SC-18 HiFi
Coverage
SC-18 assembly
Genes
IGHV3-43D
IGHV3-38-3
IGHV1-38-4
IGHV3-43


b

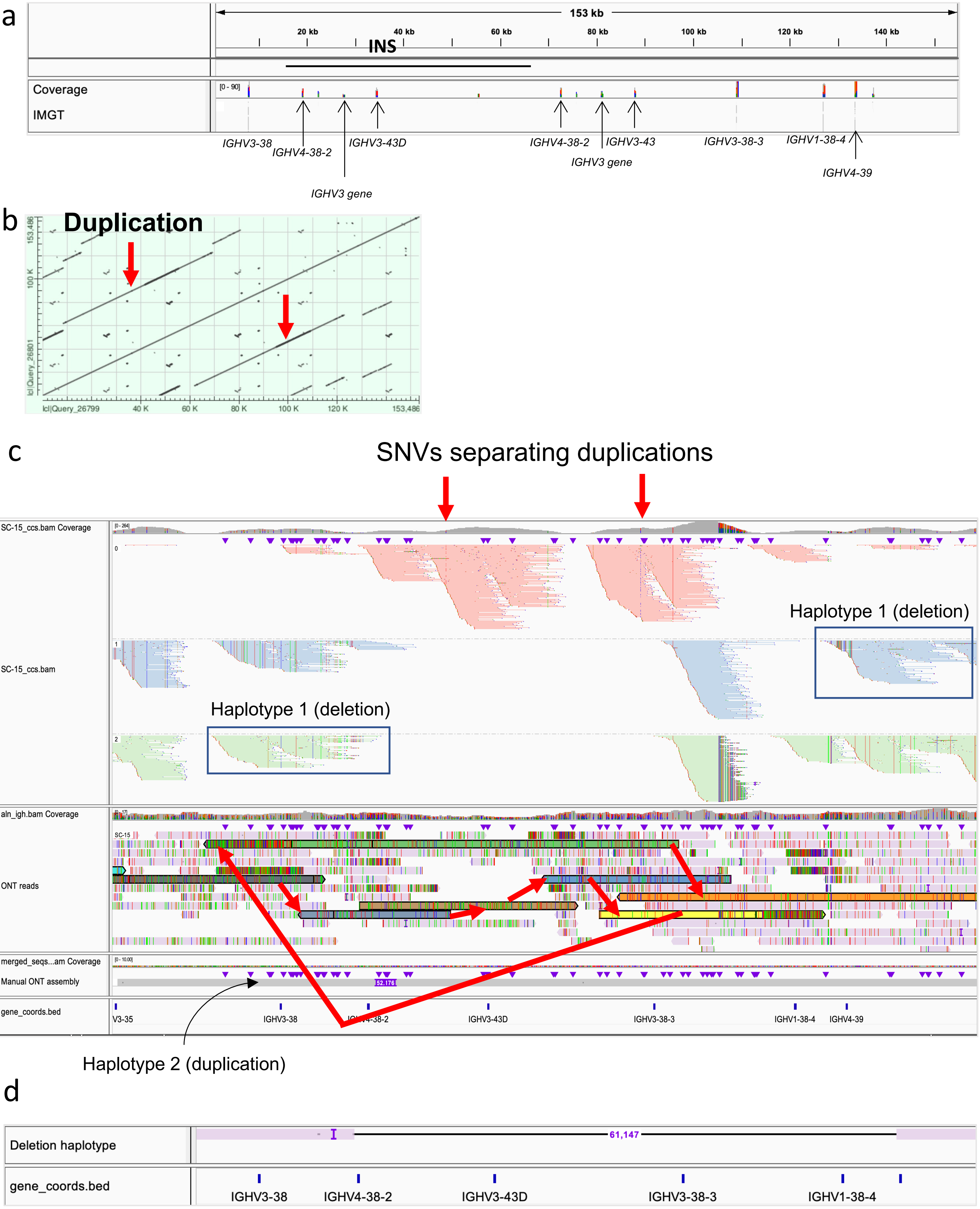
a
153 kb
20 kb
40 kb
60 kb
80 kb
100 kb
120 kb
140 kb
INS
Coverage
IMGT
[0 - 90]
IGHV3-38
IGHV4-38-2
IGHV3-43D
IGHV3 gene
IGHV4-38-2
IGHV3-43
IGHV3 gene
IGHV3-38-3
IGHV1-38-4
IGHV4-39
b
Duplication
lcl|Query_26801
lcl|Query_26799
40 K
60 K
80 K
100 K
120 K
153,486
c
SNVs separating duplications
SC-15_ccs.bam Coverage
SC-15_ccs.bam
Haplotype 1 (deletion)
Haplotype 1 (deletion)
aln_igh.bam Coverage
ONT reads
merged_seqs...am Coverage
Manual ONT assembly
gene_coords.bed
V3-35
IGHV3-38
IGHV3-43D
IGHV3-38-3
IGHV1-38-4
IGHV4-39
Haplotype 2 (duplication)
d
Deletion haplotype
61,147
gene_coords.bed
IGHV3-38
IGHV4-38-2
IGHV3-43D
IGHV3-38-3
IGHV1-38-4

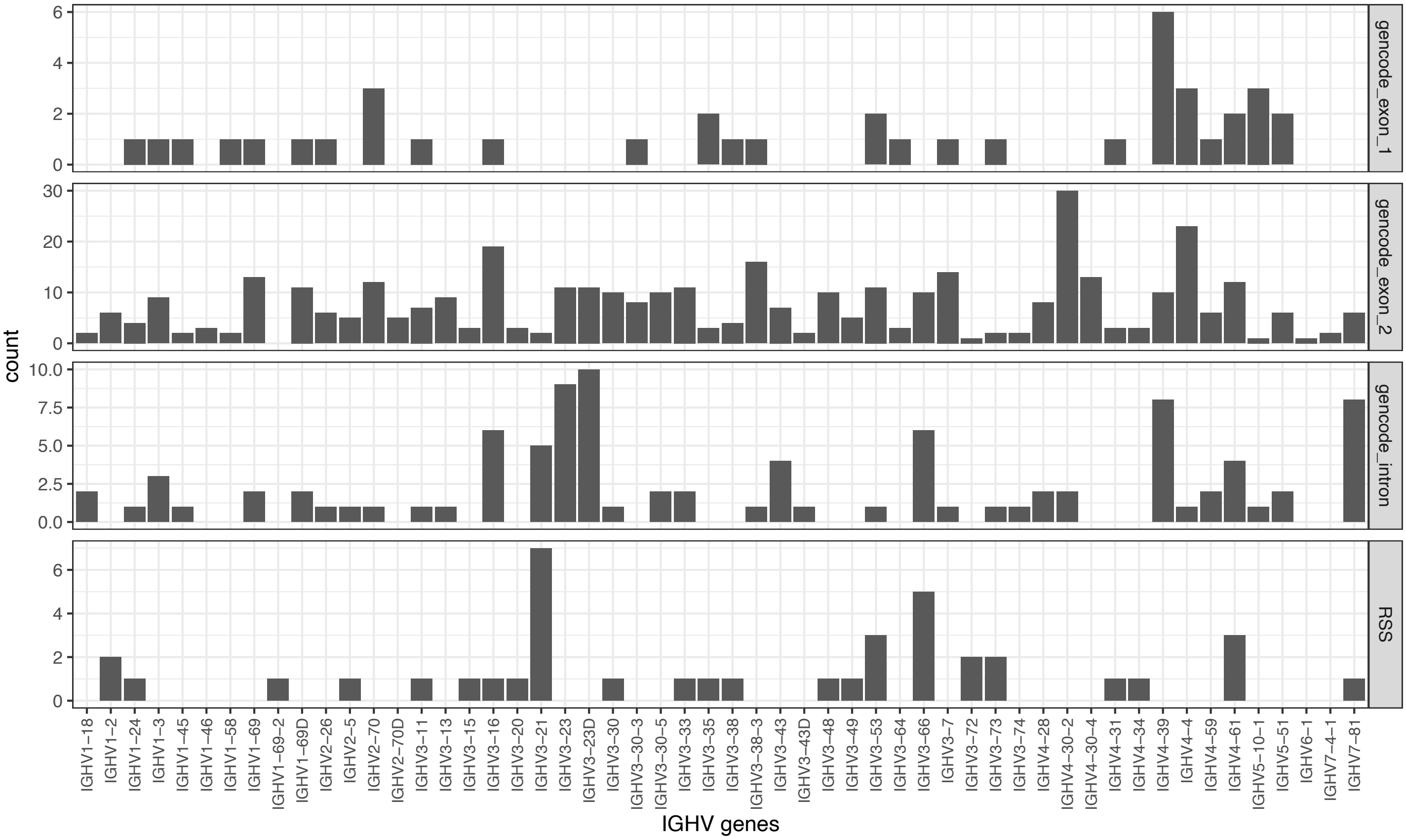

count
IGHV genes
gencode_exon_1
gencode_exon_2
gencode_intron
RSS
IGHV1−18
IGHV1−2
IGHV1−24
IGHV1−3
IGHV1−45
IGHV1−46
IGHV1−58
IGHV1−69
IGHV1−69−2
IGHV1−69D
IGHV2−26
IGHV2−5
IGHV2−70
IGHV2−70D
IGHV3−11
IGHV3−13
IGHV3−15
IGHV3−16
IGHV3−20
IGHV3−21
IGHV3−23
IGHV3−23D
IGHV3−30
IGHV3−30−3
IGHV3−30−5
IGHV3−33
IGHV3−35
IGHV3−38
IGHV3−38−3
IGHV3−43
IGHV3−43D
IGHV3−48
IGHV3−49
IGHV3−53
IGHV3−64
IGHV3−66
IGHV3−7
IGHV3−72
IGHV3−73
IGHV3−74
IGHV4−28
IGHV4−30−2
IGHV4−30−4
IGHV4−31
IGHV4−34
IGHV4−39
IGHV4−4
IGHV4−59
IGHV4−61
IGHV5−10−1
IGHV5−51
IGHV6−1
IGHV7−4−1
IGHV7−81

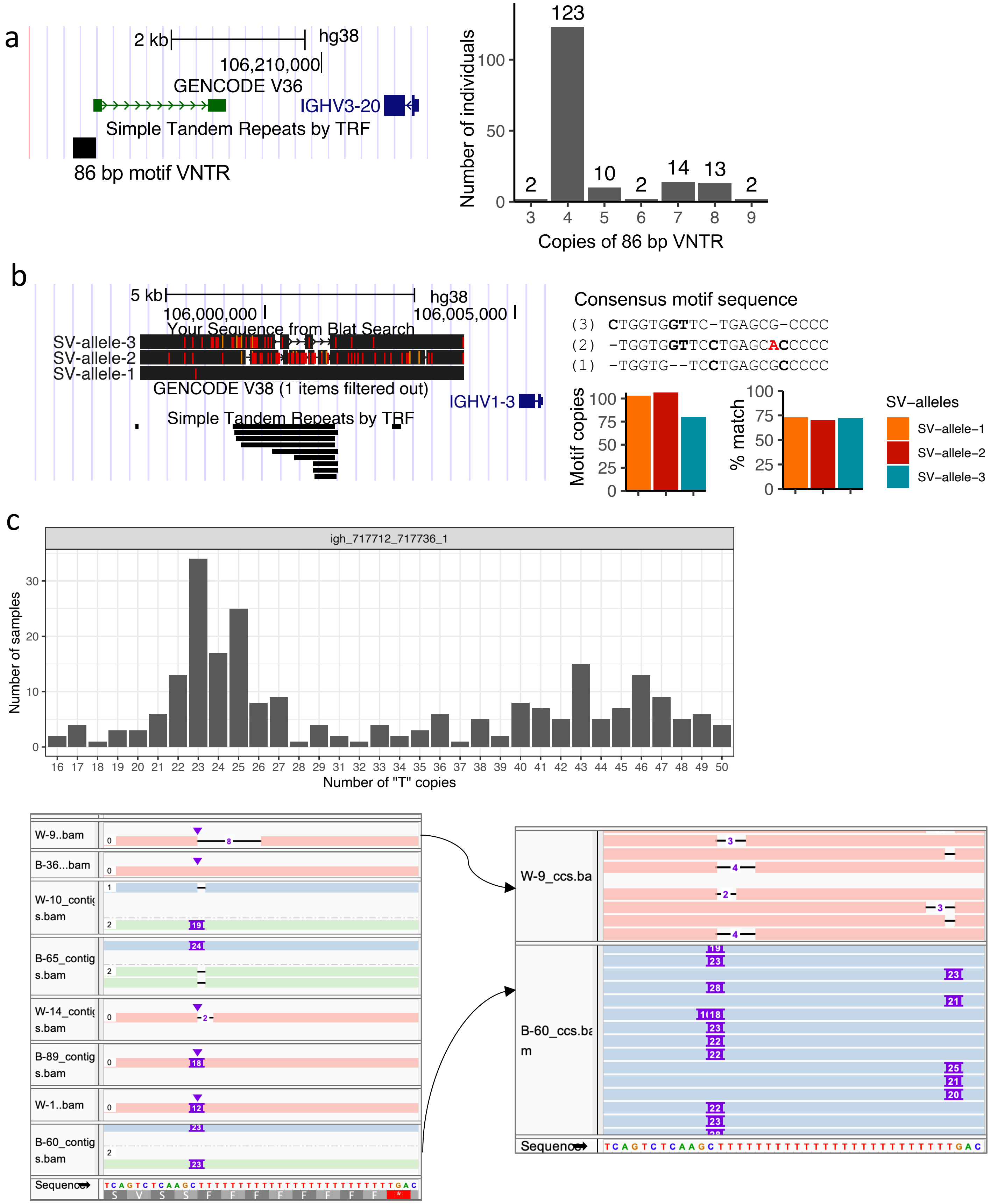
a
2 kb
hg38
106,210,000
GENCODE V36
IGHV3-20
Simple Tandem Repeats by TRF
86 bp motif VNTR
Number of individuals
Copies of 86 bp VNTR
123
2
10
2
14
13
2
b
5 kb
hg38
106,000,000
106,005,000
Your Sequence from Blat Search
SV-allele-3
SV-allele-2
SV-allele-1
GENCODE V38 (1 items filtered out)
IGHV1-3
Simple Tandem Repeats by TRF
Consensus motif sequence
(3) CTGGTGGTTC-TGAGCG-CCCC
(2) -TGGTGGTTCCTGAGCACCCCC
(1) -TGGTG--TCCTGAGCGCCCCC
Motif copies
% match
SV–alleles
SV–allele–1
SV–allele–2
SV–allele–3
c
igh_717712_717736_1
Number of samples
Number of "T" copies
W-9..bam
B-36...bam
W-10_contig s.bam
B-65_contig s.bam
W-14_contig s.bam
B-89_contig s.bam
W-1..bam
B-60_contig s.bam
Sequence
W-9_ccs.ba
B-60_ccs.bam
Sequence

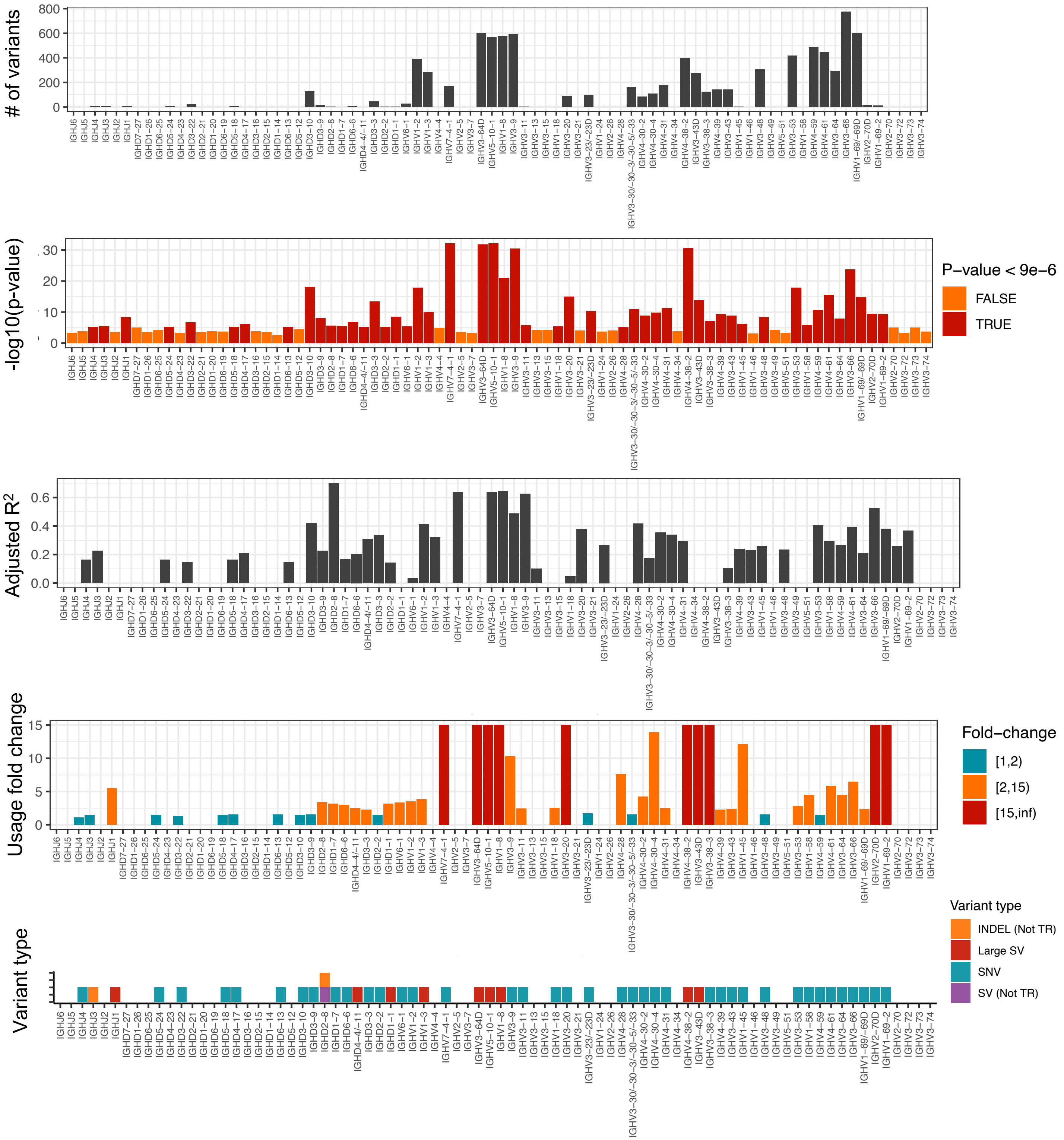

### of variants
0
200
400
600
800
-log10(p-value)
0
10
20
30
P−value < 9e−6
FALSE
TRUE
Adjusted R²
0.0
0.2
0.4
0.6
Usage fold change
0
5
10
15
Fold−change
[1,2)
[2,15)
[15,inf)
Variant type
INDEL (Not TR)
Large SV
SNV
SV (Not TR)
IGHJ6
IGHJ5
IGHJ4
IGHJ3
IGHJ2
IGHJ1
IGHD7−27
IGHD1−26
IGHD6−25
IGHD5−24
IGHD4−23
IGHD3−22
IGHD2−21
IGHD1−20
IGHD6−19
IGHD5−18
IGHD4−17
IGHD3−16
IGHD2−15
IGHD1−14
IGHD6−13
IGHD5−12
IGHD3−10
IGHD3−9
IGHD2−8
IGHD1−7
IGHD6−6
IGHD4−4/−11
IGHD3−3
IGHD2−2
IGHD1−1
IGHV6−1
IGHV1−2
IGHV1−3
IGHV4−4
IGHV7−4−1
IGHV2−5
IGHV3−7
IGHV3−64D
IGHV5−10−1
IGHV1−8
IGHV3−9
IGHV3−11
IGHV3−13
IGHV3−15
IGHV1−18
IGHV3−20
IGHV3−21
IGHV3−23/−23D
IGHV1−24
IGHV2−26
IGHV4−28
IGHV3−30/−3/−30−3/−30−5/−33
IGHV4−30−2
IGHV4−30−4
IGHV4−31
IGHV4−34
IGHV4−38−2
IGHV3−43D
IGHV3−38−3
IGHV4−39
IGHV3−43
IGHV1−45
IGHV1−46
IGHV3−48
IGHV3−49
IGHV5−51
IGHV3−53
IGHV1−58
IGHV4−59
IGHV4−61
IGHV3−64
IGHV3−66
IGHV1−69/−69D
IGHV2−70D
IGHV1−69−2
IGHV2−70
IGHV3−72
IGHV3−73
IGHV3−74

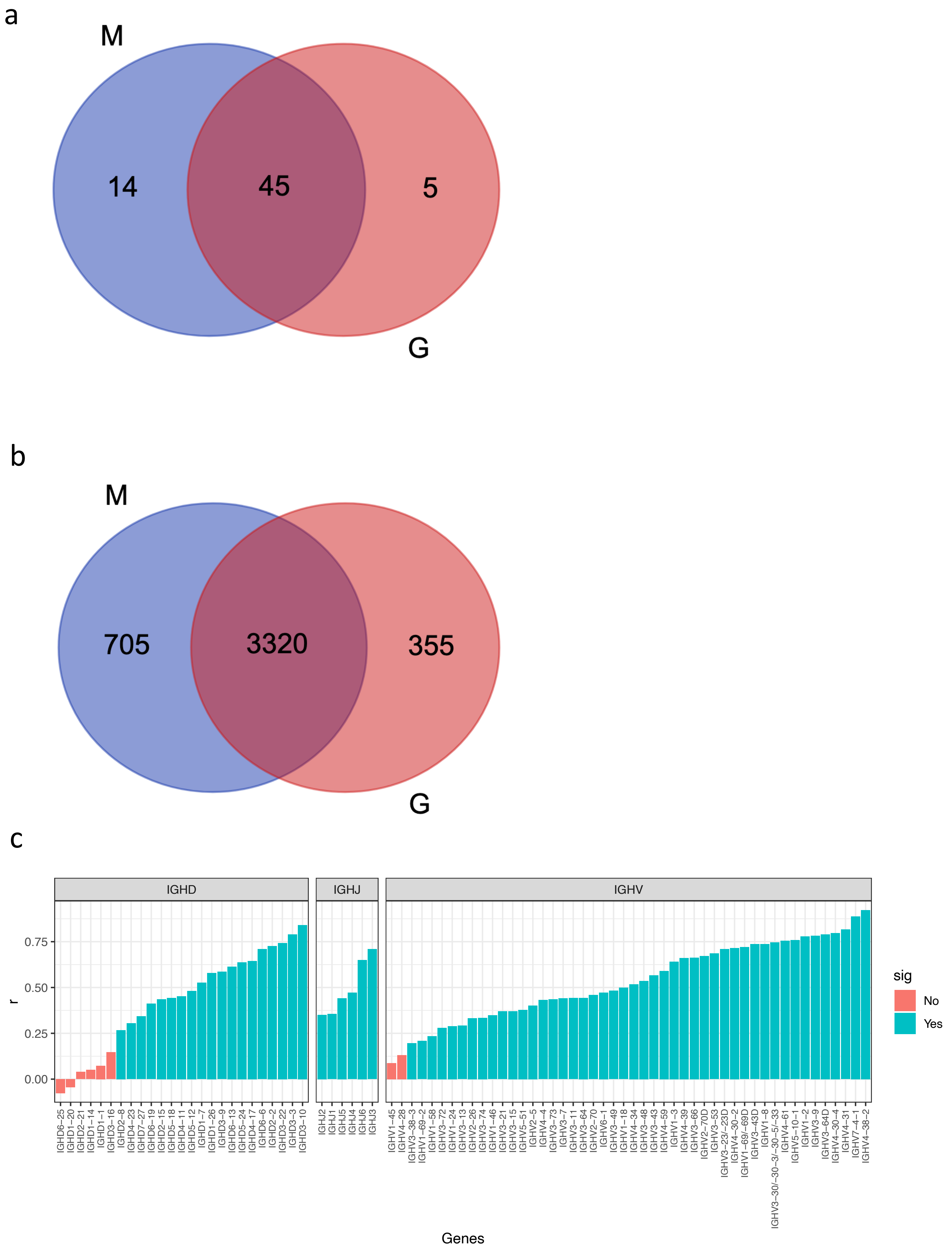
a
M
G
14
45
5
b
M
G
705
3320
355
c
IGHD
IGHJ
IGHV
r
0.00
0.25
0.50
0.75
sig
No
Yes
Genes
IGHD6−25
IGHD1−20
IGHD2−21
IGHD1−14
IGHD1−1
IGHD3−16
IGHD2−8
IGHD4−23
IGHD7−27
IGHD6−19
IGHD2−15
IGHD5−18
IGHD4−11
IGHD5−12
IGHD1−7
IGHD1−26
IGHD3−9
IGHD6−13
IGHD5−24
IGHD4−17
IGHD6−6
IGHD2−2
IGHD3−22
IGHD3−3
IGHD3−10
IGHJ2
IGHJ1
IGHJ5
IGHJ4
IGHJ6
IGHJ3
IGHV1−45
IGHV4−28
IGHV3−38−3
IGHV1−69−2
IGHV1−58
IGHV3−72
IGHV1−24
IGHV3−13
IGHV2−26
IGHV3−74
IGHV1−46
IGHV3−21
IGHV3−15
IGHV5−51
IGHV2−5
IGHV4−4
IGHV3−73
IGHV3−7
IGHV3−11
IGHV3−64
IGHV2−70
IGHV6−1
IGHV3−49
IGHV1−18
IGHV4−34
IGHV3−48
IGHV3−43
IGHV4−59
IGHV1−3
IGHV4−39
IGHV3−66
IGHV2−70D
IGHV3−53
IGHV3−23/−23D
IGHV4−30−2
IGHV1−69/−69D
IGHV3−43D
IGHV1−8
IGHV3−30/−30−3/−30−5/−33
IGHV4−61
IGHV5−10−1
IGHV1−2
IGHV3−9
IGHV3−64D
IGHV4−30−4
IGHV4−31
IGHV7−4−1
IGHV4−38−2

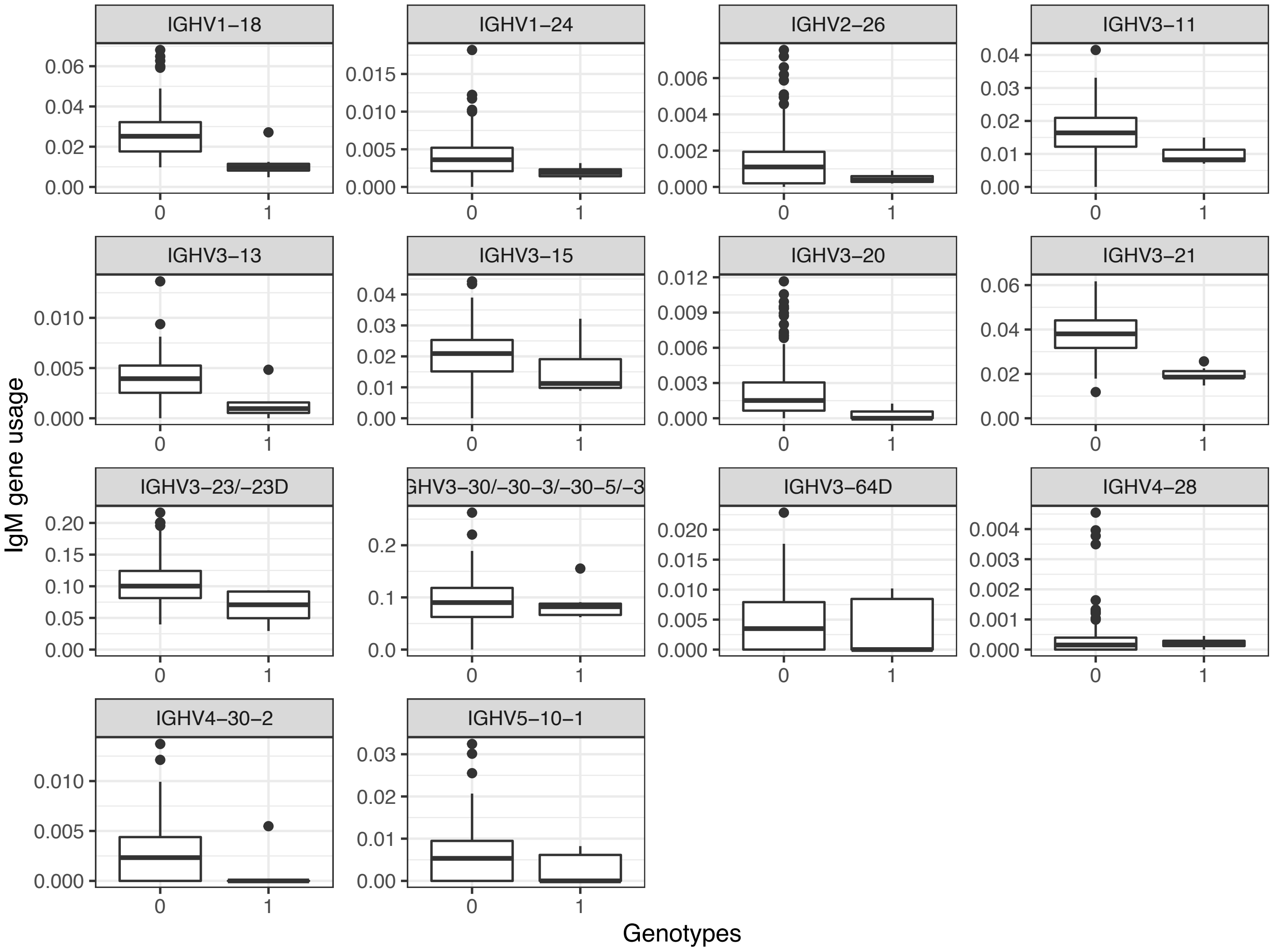
IGHV1−18
IGHV1−24
IGHV2−26
IGHV3−11
IGHV3−13
IGHV3−15
IGHV3−20
IGHV3−21
IGHV3−23/−23D
GHV3−30/−30−3/−30−5/−3
IGHV3−64D
IGHV4−28
IGHV4−30−2
IGHV5−10−1
IgM gene usage
Genotypes

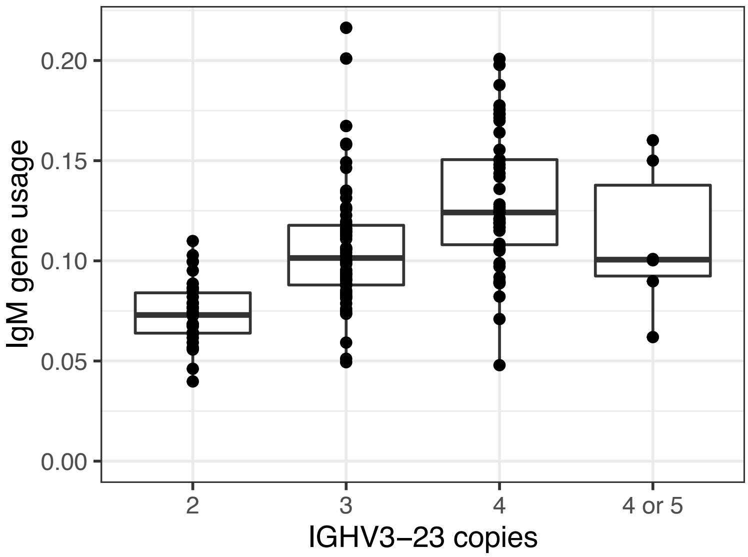

0.20
0.15
0.10
0.05
0.00
IgM gene usage
2
3
4
4 or 5
IGHV3−23 copies

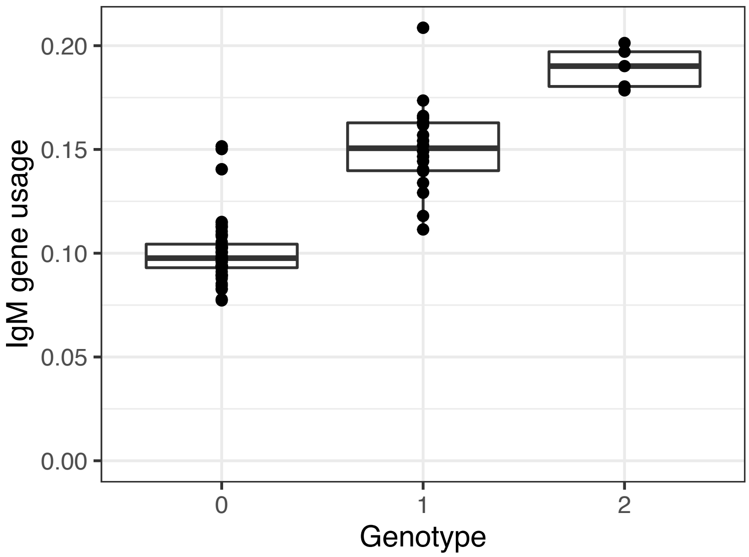

IgM gene usage
0.20
0.15
0.10
0.05
0.00
0
1
2
Genotype

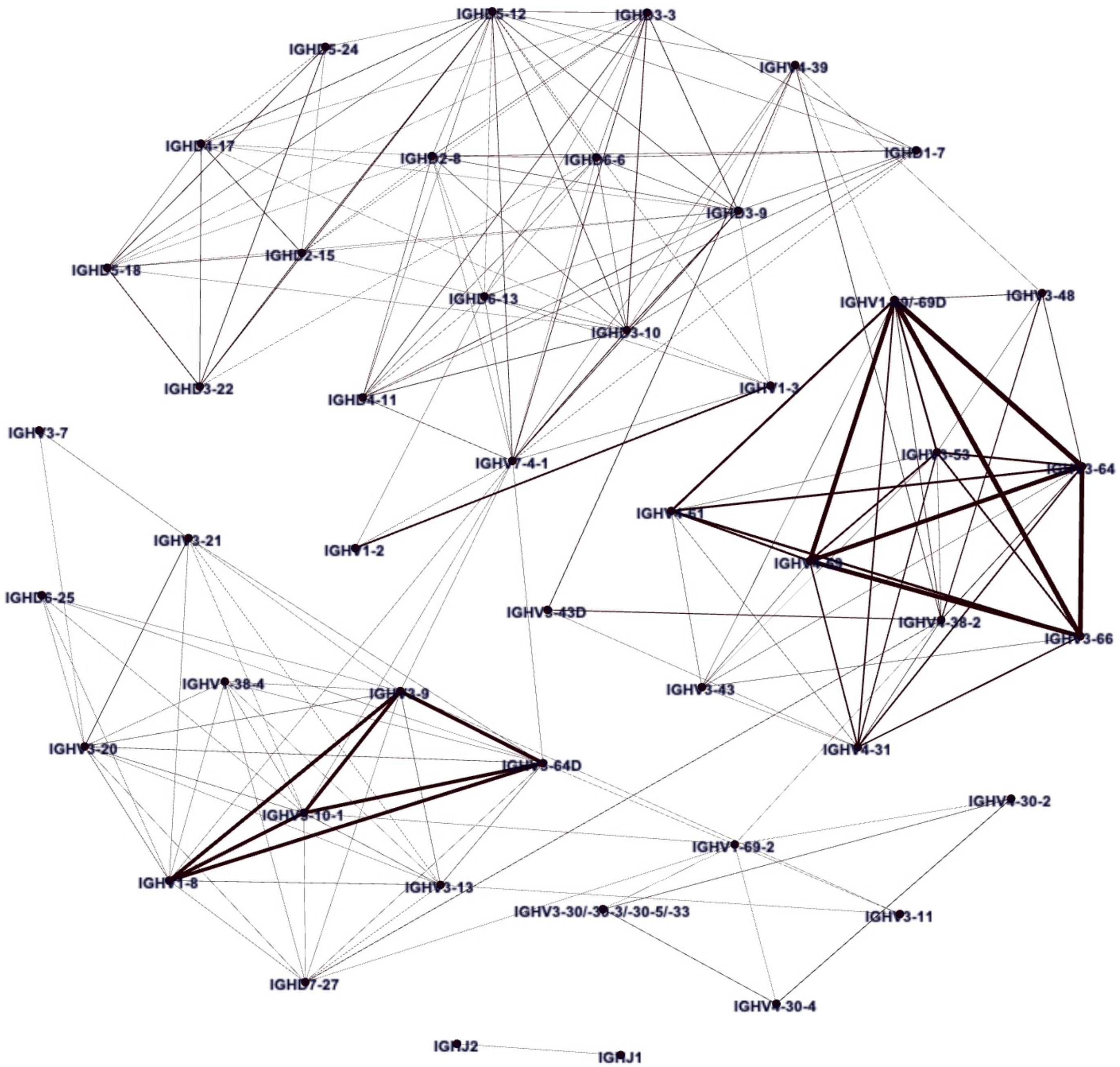
IGHD5-12
IGHD3-3
IGHD5-24
IGHV4-39
IGHD4-17
IGHD2-8
IGHD6-6
IGHD1-7
IGHD3-9
IGHD2-15
IGHD5-18
IGHD6-13
IGHV1-69/-69D
IGHV3-48
IGHD3-10
IGHD3-22
IGHD4-11
IGHV1-3
IGHV3-7
IGHV7-4-1
IGHV3-53
IGHV3-64
IGHV4-61
IGHV3-21
IGHV1-2
IGHV4-59
IGHD6-25
IGHV3-43D
IGHV4-38-2
IGHV3-66
IGHV1-38-4
IGHV3-43
IGHV3-9
IGHV3-20
IGHV3-64D
IGHV4-31
IGHV4-30-2
IGHV5-10-1
IGHV1-69-2
IGHV1-8
IGHV3-13
IGHV3-30/-30-3/-30-5/-33
IGHV3-11
IGHD7-27
IGHV4-30-4
IGHJ2
IGHJ1

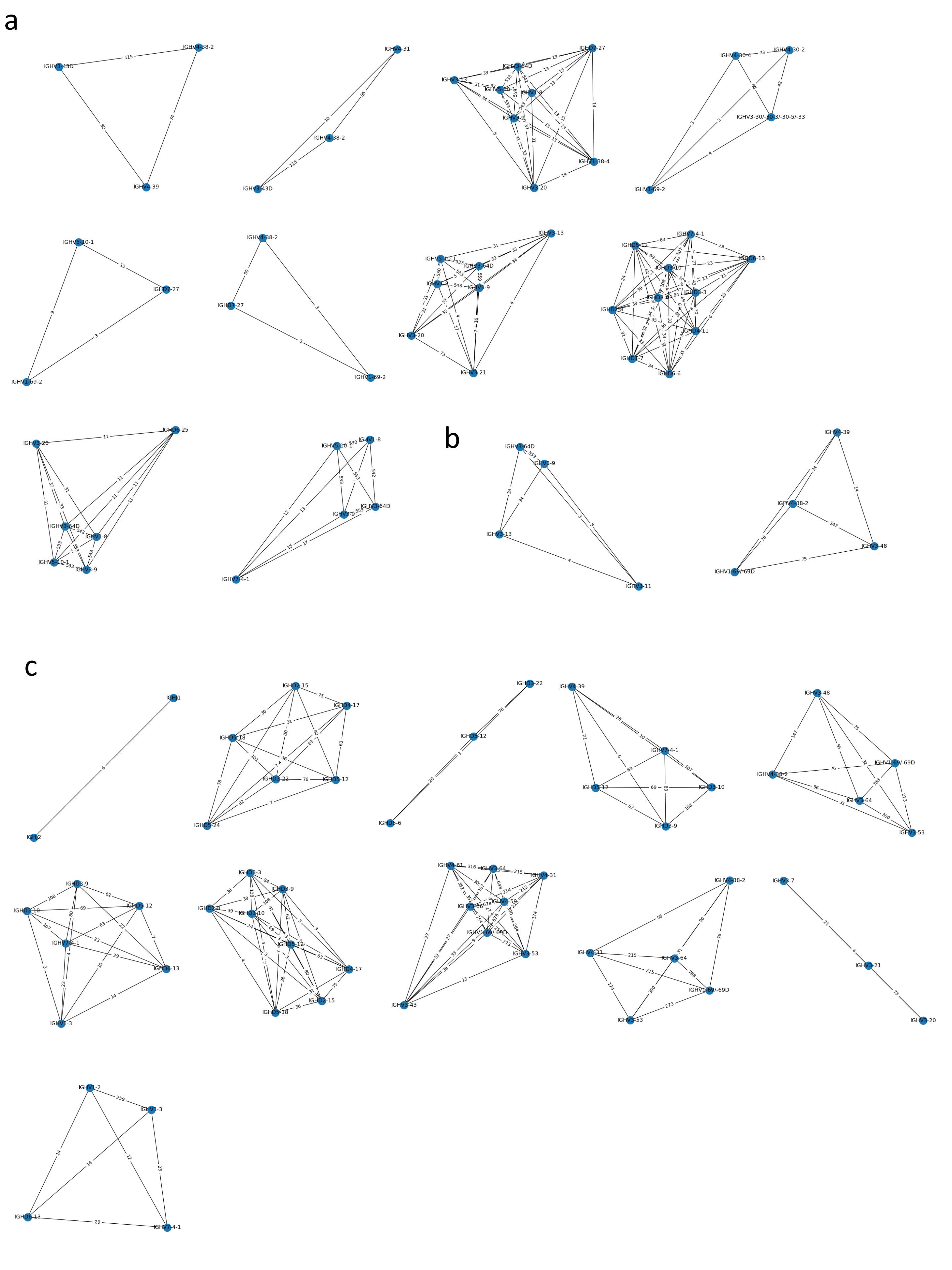
a
IGHV4-38-2
IGHV3-43D
IGHV4-39
IGHV4-31
IGHV4-38-2
IGHV3-43D
IGHV7-27
IGHV3-64D
IGHV3-13
IGHV1-8
IGHV3-9
IGHV1-38-4
IGHV3-20
IGHV4-30-4
IGHV4-30-2
IGHV3-30/-30-3/-30-5/-33
IGHV1-69-2
IGHV5-10-1
IGHV7-27
IGHV1-69-2
IGHV4-38-2
IGHV7-27
IGHV1-69-2
IGHV3-13
IGHV5-10-1
IGHV3-64D
IGHV1-8
IGHV3-9
IGHV3-20
IGHV3-21
IGHD5-12
IGHV7-4-1
IGHD6-13
IGHD3-10
IGHD3-3
IGHD2-8
IGHD4-11
IGHD1-7
IGHD6-6
IGHV3-20
IGHD6-25
IGHV3-64D
IGHV1-8
IGHV5-10-1
IGHV3-9
IGHV5-10-1
IGHV1-8
IGHV3-9
IGHV3-64D
IGHV7-4-1
b
IGHV3-64D
IGHV3-9
IGHV3-13
IGHV3-11
IGHV4-39
IGHV4-38-2
IGHV3-48
IGHV1-69/-69D
c
IGHJ1
IGHJ2
IGHD2-15
IGHD4-17
IGHD5-18
IGHD3-22
IGHD5-12
IGHD5-24
IGHD3-22
IGHD5-12
IGHD6-6
IGHV4-39
IGHV7-4-1
IGHD5-12
IGHD3-10
IGHD3-9
IGHV3-48
IGHV1-69/-69D
IGHV4-38-2
IGHV3-64
IGHV3-53
IGHD3-9
IGHD3-10
IGHD5-12
IGHV7-4-1
IGHD6-13
IGHV1-3
IGHD3-3
IGHD3-9
IGHD2-8
IGHD3-10
IGHD5-12
IGHD4-17
IGHD5-18
IGHD2-15
IGHV4-61
IGHV3-64
IGHV4-31
IGHV3-66
IGHV4-59
IGHV1-69/-69D
IGHV3-53
IGHV3-43
IGHV4-38-2
IGHV4-31
IGHV3-64
IGHV1-69/-69D
IGHV3-53
IGHV3-7
IGHV3-21
IGHV3-20
IGHV1-2
IGHV1-3
IGHD6-13
IGHV7-4-1
